# Low-Dimensional Organization of Metabolic Phenotypes Underlies Recurrent Evolutionary Diversification

**DOI:** 10.64898/2026.04.22.720058

**Authors:** Keita Saito, Saburo Tsuru, Chikara Furusawa

## Abstract

Metabolic phenotypes vary within microbial species, yet how such variation is organized remains unclear. Diversification in carbon-source utilization often appears idiosyncratic, showing weak correspondence to phylogeny or simple gene content. Here, we combine quantitative growth phenotyping of natural *Escherichia coli* isolates across 32 carbon sources with diversification observed during de novo laboratory evolution, together with a reaction-level description of metabolic similarity. Despite deep phylogenetic divergence, growth-rate profiles varied independently of lineage. Instead, growth rates across carbon sources covaried in recurrent modular patterns aligned with flux-based metabolic similarity. Closely analogous modular relationships re-emerged during de novo evolution, indicating parallel diversification across evolutionary contexts. Growth-rate variation in natural and experimentally evolved datasets collapsed onto a shared low-dimensional variance structure. Together, our results indicate that quantitative metabolic phenotypes vary along a limited set of recurring, module-linked axes, providing insight into why similar metabolic states can repeatedly arise across independent evolutionary contexts.

## Introduction

Biological traits rarely diversify at random. Instead, the directions along which phenotypes vary and evolve are shaped not only by natural selection but also by how phenotypic variability is distributed across traits and their combinations^1^. Because underlying variation is unevenly distributed across these combinations^2–6^, some phenotypic changes respond readily to selection whereas others change only slowly ^7,8^, resulting in biased—not random—trajectories through phenotypic space^9–11^. Understanding how such biases in phenotypic variation arise is a central goal in evolutionary biology, as it determines which forms of phenotypic diversity are accessible during evolution.

A prominent source of such organization is the architecture of biological networks^11^. Developmental and regulatory networks can couple otherwise distinct traits, concentrate variation along a few shared directions rather than independently, and thereby generate correlated responses^12,13^. Gene regulatory networks provide a clear example of this effect: across many systems, they have been shown to concentrate variation in expression levels^4,5^ into a low-dimensional structure, leading to correlated patterns of phenotypic variation^5,12^. From this viewpoint, evolutionary diversification is expected to reflect, at least in part, the internal organization of the networks that generate phenotypes^11^.

Metabolism provides a natural and experimentally tractable system in which to examine these ideas. Metabolic phenotypes arise from interconnected reaction networks governed by stoichiometric constraints and have been reported to exhibit hierarchical and modular organization^14–16^. Consistent with this view, genome-scale metabolic models and flux balance analysis have shown that metabolic network structure can account for broad patterns in growth and flux allocation^17–19^, suggesting that metabolic architecture can bias the directions along which growth phenotypes vary. At the same time, microbial metabolism presents a particularly challenging test case. Metabolism lies at the core of cellular physiology and ecological interactions, yet metabolic traits often vary strikingly even among closely related strains.

Empirical studies of natural isolates underscore this difficulty. Surveys of *Escherichia coli* have reported extensive variation in growth across carbon sources, often with little correspondence to phylogeny or to the simple presence or absence of metabolic pathways^20^. For example, Sabarly et al.^20^ showed that multilocus phylogeny was a poor predictor of quantitative metabolic phenotypes, and that while gene content could account for average growth capabilities, it failed to explain growth differences among strains. Experimental evolution studies have likewise provided limited evidence for coordinated diversification across carbon sources when evaluated under designs not optimized for this question^21,22^. Consequently, metabolic diversification within species has often been interpreted as largely idiosyncratic.

This apparent unpredictability within species stands in stark contrast to patterns observed at broader phylogenetic scales. Large-scale comparative studies across diverse bacterial species have shown that interspecific metabolic preferences often collapse onto a few major axes—most notably a trade-off between sugar and organic-acid utilization—that are broadly predictable from genomic attributes such as pathway gene counts or GC content ^23^. A complementary view emerges from community ecology: Vila et al. ^24^ found clear phylogenetic signal at the family level in carbon-source utilization among Proteobacteria, with fermentative and respiratory clades differing systematically in growth preferences across glycolytic versus gluconeogenic substrates ^24^. Yet this family-level structure coexisted with substantial quantitative variation in community composition among strains within each family. Together, these findings suggest that macro-evolutionary axes captured by genomic attributes, while powerful across broad taxonomic scales, leave much of the quantitative metabolic variation among closely related strains unaccounted for.

Importantly, this limitation does not imply that intraspecific metabolic diversification is unstructured in the directions along which phenotypes vary. Rather, these findings point to a gap between descriptions based on simple gene- or pathway-level presence–absence or GC content and our ability to assess whether quantitative phenotypic variation across metabolic conditions within a species—where genomes are largely shared—is structured along specific directions. Without a framework that links quantitative phenotypic variation to similarities among carbon sources at the level of metabolic reactions, it has remained unclear whether such apparent unpredictability reflects truly unconstrained variation or instead masks structured, directional organization rooted in metabolic network architecture.

Here, we combine quantitative growth phenotyping of natural *E. coli* isolates across diverse carbon sources with diversification observed during de novo laboratory evolution, together with genome-scale metabolic modeling to quantify distances among carbon sources. Across both natural isolates and laboratory-evolved populations, we find that growth rates do not vary independently across carbon sources, but instead covary along recurrent directions, revealing modular patterns of metabolic diversification. These directional patterns are largely independent of phylogenetic relationships, yet align with these metabolic distances. Strikingly, closely analogous patterns re-emerge during de novo evolution, and growth-rate variation in both natural and experimentally evolved datasets collapses onto a shared low-dimensional structure. Together, our results indicate that quantitative metabolic phenotypes vary along a limited set of recurring, module-linked axes, providing insight into why similar metabolic phenotypes repeatedly emerge across independent evolutionary histories.

## Results

### Growth rates of natural isolates of *E. coli* among carbon sources

We quantified carbon-source utilization profiles for 72 natural isolates of *E. coli* belonging to the ECOR collection^25^, which spans the major phylogroups^26^ (A, B1, B2, C, D, E, and F) (**Figure 1A**, **Table S1**). Growth was assayed on 32 carbon sources covering diverse biochemical classes (monosaccharides, disaccharides, amino sugars, sugar alcohols, carboxylic acids, and amino acids) and encompassing distinct entry points into central metabolism (**Figure 1B, Table S2**). Four biological replicates were measured per strain–carbon combination using a 384-well microplate (**Table S3**). In total, 9216 cultures were monitored for 72 hours using a high-throughput automated system^27^ to obtain growth curves (**Figure 1C, Figure S1**). From these growth curves, we estimated the maximal growth rate (μ_max_) for each culture using a logistic growth model. The average growth rate across replicates for each strain–carbon combination was used for downstream analysis (**Figures S2** and **S3**; **Table S4**).

**Figure 1.**
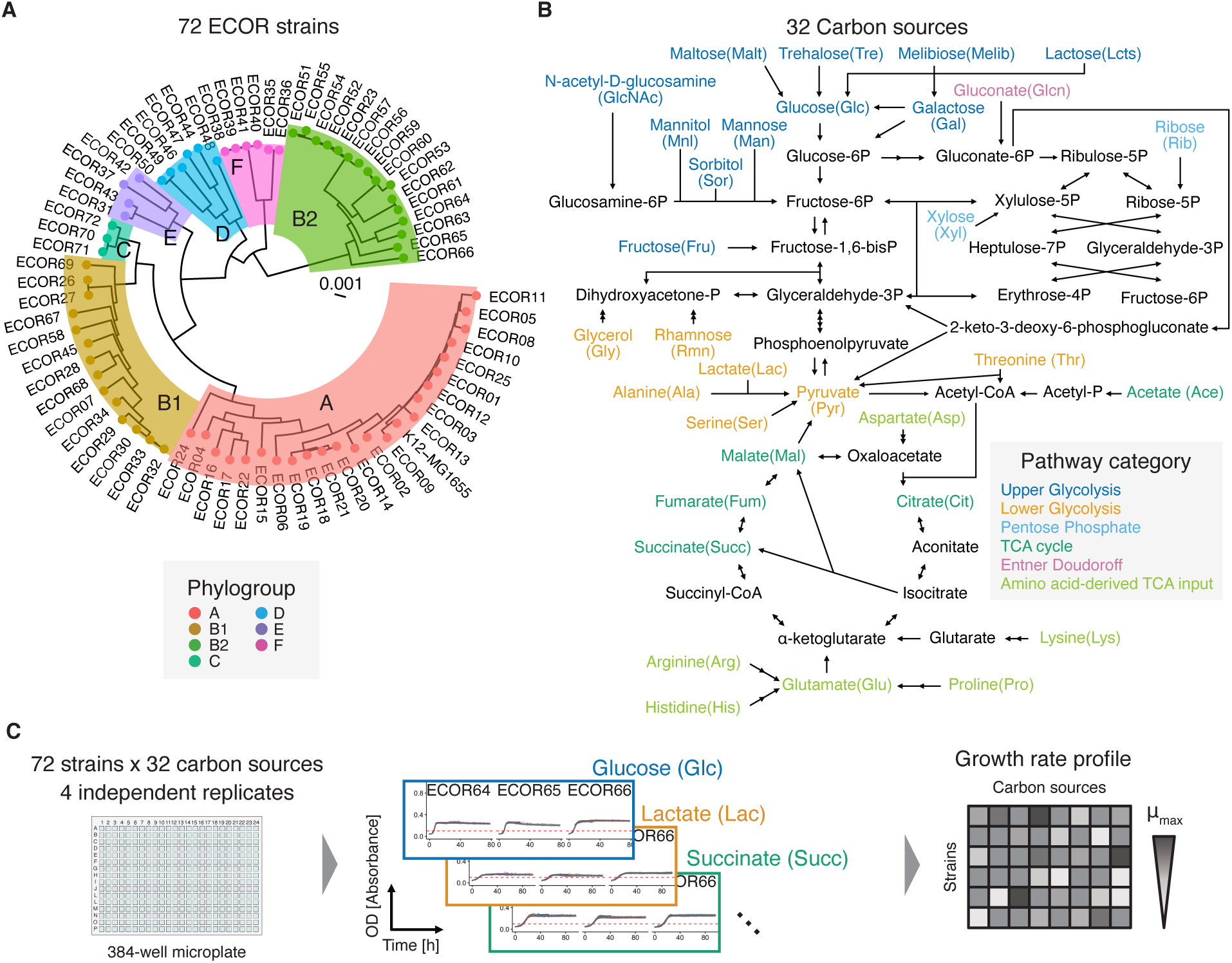
Experimental overview for quantifying growth-rate profiles across *E. coli* strains and carbon sources. (**A**) Maximum-likelihood phylogenetic tree of 72 ECOR strains and the K-12 MG1655 strain. (**B**) Simplified schematic of the *E. coli* metabolic network. Carbon sources used in this study are colored according to their metabolic pathway categories. (**C**), Experimental workflow. Combinations of 72 strains and 32 carbon sources were cultured in microplates (left). Optical density (OD) was monitored at regular intervals using an automated culture system. Growth curves were analyzed to estimate maximum growth rate (μ_max_) (middle). The resulting growth-rate profiles were used for downstream analyses (right).

### Decoupling between phylogenetic distance and diversification in carbon-source utilization

To examine evolutionary patterns in growth-rate profiles, we first tested whether closely related strains exhibit more similar profiles. For each strain pair, we calculated Spearman’s rank correlation (R) of μ_max_ across carbon sources, as a measure of growth similarity (**Figure 2A**). Some closely related strains such as ECOR01 and ECOR02 showed a strong correlation (Spearman’s R = 0.89, P = 2.2 × 10^−6^, **Figure 2B**), whereas some distantly related strains such as ECOR01 and ECOR50 showed weaker correlations (Spearman’s R = 0.55, P = 5.1 × 10^−3^, **Figure 2C**). Hierarchical clustering of pairwise correlations generated a dendrogram representing similarity in growth-rate profiles across strains (**Figure 2D**, bottom). The corresponding tanglegram revealed a moderate but statistically non-significant topological agreement between the phylogeny and the growth-based dendrogram (entanglement score = 0.32, P = 0.17), indicating that divergence in growth-rate profiles only weakly or partially reflects phylogenetic branching order. Indeed, major phylogroups did not form distinct clusters in the growth-based dendrogram, and distantly related strains frequently exhibited highly similar profiles. This weak structural congruence was also confirmed topologically (Baker’s gamma = 0.041, P = 0.034).

**Figure 2.**
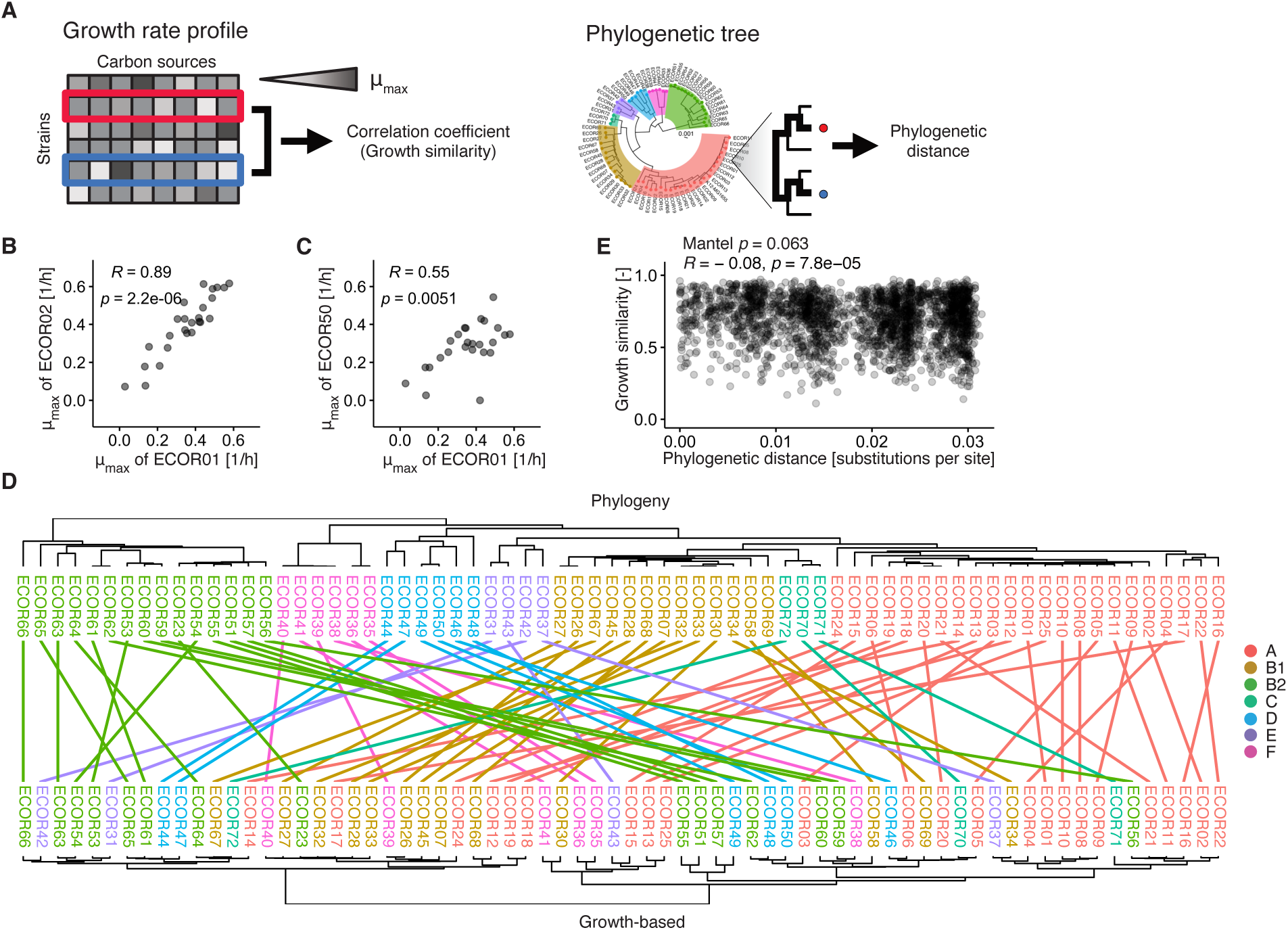
Decoupling between phylogeny and growth-rate profiles. (**A**) Schematic representation of pairwise comparisons across strains. Growth similarity between strains was quantified using Spearman’s rank correlation (R) in μ_max_ (left). Phylogenetic distance was defined as pairwise patristic distance on the phylogenetic tree (right). (**B**) and (**C**) Representative examples showing relationships in μ_max_ between strain pairs: a closely related pair (ECOR01 and ECOR02, **B**) and a distantly related pair (ECOR01 and ECOR50, **C**). (**D**) Tanglegram comparing the phylogenetic tree (top) and the growth-based dendrogram (bottom). Lines connect the same strains across trees. The entanglement score was 0.32. (**E**) Relationship between growth similarity and phylogenetic distance across all strain pairs. P-value in Mantel test with 999 permutations is indicated; the Spearman P-value shown in the inset is for reference only, as pairwise distance matrices violate the independence assumption of standard correlation tests. Insets in panels (**B**), **(C**), and (**E**) show Spearman’s rank correlation coefficients (R) and associated two-sided P-values.

To incorporate evolutionary time, we next examined the relationship between patristic distance on the ECOR maximum-likelihood tree (**Figure 1A**), and the similarity in growth-rate profiles across strain pairs. Mantel test ^28^ revealed no significant association between patristic distance and growth similarity (Mantel test, P = 0.063; Spearman’s R = −0.08, **Figure 2E**). Consistent with this, Blomberg’s K ^29^ was extremely low across all carbon sources (**Figure S4**), indicating negligible phylogenetic signal. Together, these findings demonstrate that diversification in carbon-source utilization is largely phylogenetically decoupled, suggesting that growth-rate profiles are not constrained by ancestral states and may diversify rapidly in a parallel or convergent manner.

### Interstrain correlation in growth-rate profiles across carbon sources

Although growth-rate profiles diversified largely independently of phylogeny, they may nonetheless be associated with metabolic pathway architecture. To test this, we assessed interstrain correlations in growth rates across carbon sources. For each carbon-source pair, we computed Spearman’s R of μ_max_ across strains (**Figure 3A–C**). Given the negligible phylogenetic signal indicated by Blomberg’s K (**Figure S4**), the interstrain correlations were expected to predominantly reflect genuine evolutionary correlations. To confirm this expectation and to account for potential phylogenetic confounding when evaluating evolutionary correlations across carbon sources in ECOR strains, we calculated phylogenetic signal-adjusted correlations using two approaches: Phylogenetic Generalized Least Squares (PGLS) and Phylogenetically Independent Contrasts (PIC). As expected, both PGLS- and PIC-derived evolutionary correlations showed strong agreement with interstrain correlations (PGLS, Spearman’s R = 0.96, P < 2.2 × 10^−16^; Mantel test, P = 0.001; PIC, Spearman’s R = 0.95, P < 2.2 × 10 ^−16^; Mantel test, P = 0.001; **Figure S5**). For reference throughout this analysis, carbon sources were grouped into conventional metabolic pathway categories^30^ based on their primary entry points into central metabolism, including upper glycolysis, lower glycolysis, the tricarboxylic acid (TCA) cycle, the Entner–Doudoroff (ED) pathway, the pentose phosphate (PP) pathway, and amino-acid–derived inputs feeding directly into the TCA cycle. Hierarchical clustering of pairwise correlations revealed clear modular structure (**Figure 3D**). One cluster (E1) consisted mainly of upper-glycolysis carbon sources, while another (E2) consisted largely of TCA-cycle carbon sources, amino-acid-derived TCA inputs (e.g., proline), and many lower-glycolysis carbon sources (e.g., pyruvate). To validate the choice of two modules, analysis of the dendrogram’s fusion distances confirmed that merging the two primary clusters into a single group (k_c_ = 2) required a drastically larger distance compared to subsequent splits (such as at k_c_ = 3), supporting the definition of two primary modules (**Figure 3E**). Within-cluster correlations were significantly higher than between-cluster correlations (**Figure 3F**, Wilcoxon test, BH-adjusted P < 0.05).

**Figure 3.**
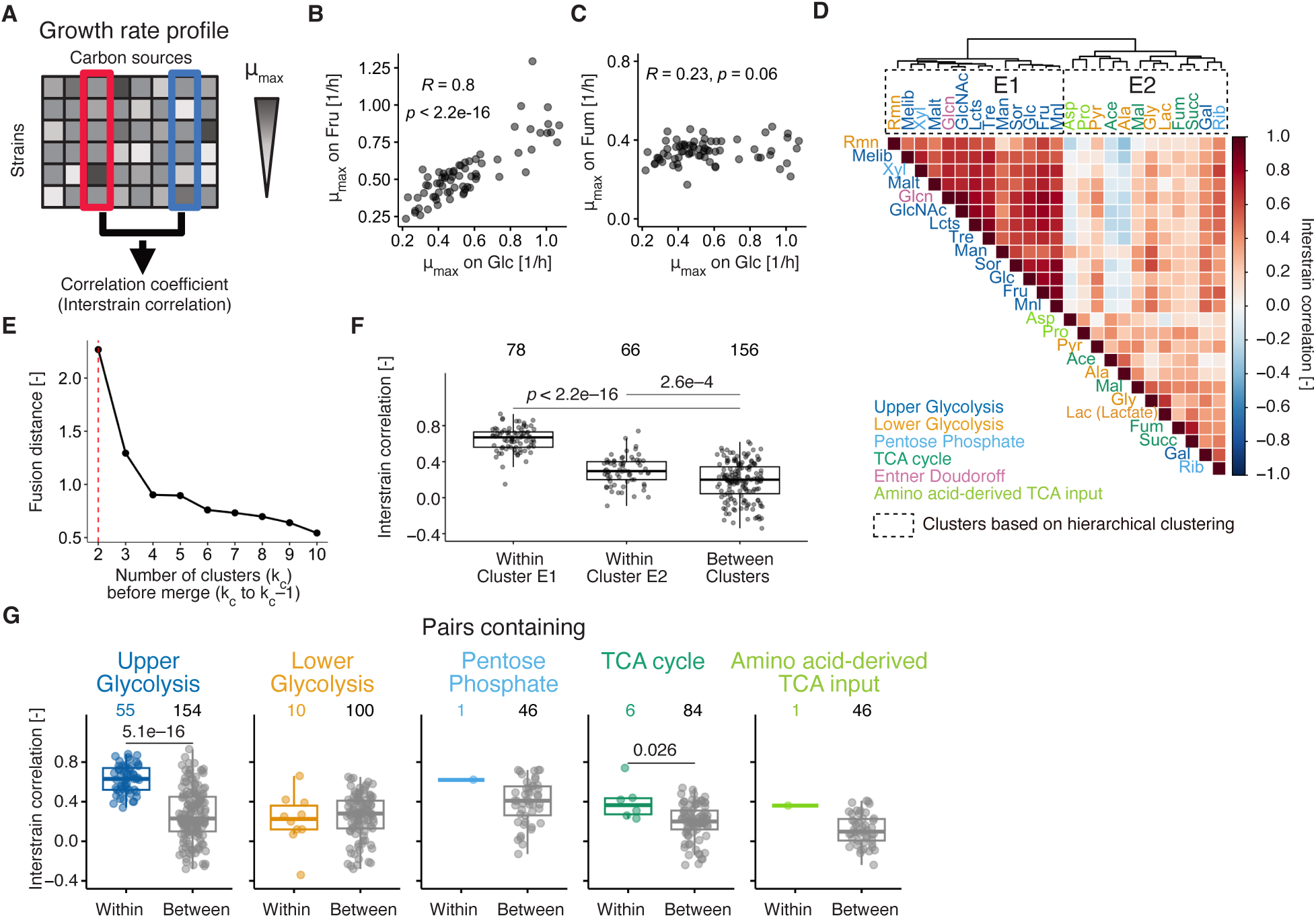
Interstrain correlations across carbon sources. (**A**) Schematic representation of pairwise analysis across carbon sources. Interstrain correlation was defined as Spearman’s rank correlation in μ_max_ across strains. (**B**) and (**C**) Representative examples of interstrain correlations: a within-category pair (**B**) and a between-category pair (**C**). Spearman’s R with P-values (two-sided) are shown. (**D**) Heatmap of interstrain correlations among all carbon-source pairs. Dashed rectangles indicate clusters (E1 and E2) identified by hierarchical clustering (Ward.D2). (**E**) Scree plot of the dendrogram’s fusion distances across different numbers of clusters (k_c_ prior to merging to k_c_ − 1). The dashed line indicates k_c_ = 2. (**F**) Comparisons of interstrain correlations for within- and between-cluster pairs for each cluster. (**G**) Comparisons of interstrain correlations for within- and between-category pairs for each metabolic pathway category. In panels (**F**) and (**G**), boxplot elements represent the median (horizontal line), first and third quartiles (q1 and q3), and whiskers extending to the most extreme data points within 1.5×(q3 − q1). P values were calculated using the Wilcoxon test (two-sided) with Benjamini-Hochberg (BH) correction. Numbers above the boxes indicate the number of pairs.

However, conventional pathway categories captured only part of this modular structure (**Figure 3G**). While strong clustering was observed for upper-glycolysis carbon sources (Wilcoxon test, BH-adjusted P = 5.1 × 10^−16^), only marginal or no significant within-category elevation of correlation was detected for TCA-cycle (Wilcoxon test, BH-adjusted P = 0.026) or lower-glycolysis carbon sources (Wilcoxon test, BH-adjusted P = 0.53). To address whether the strong significance of upper glycolysis was merely driven by its larger sample size, we performed a subsampling analysis by randomly sampling four carbon sources (matching the number of TCA-cycle substrates) from upper glycolysis over 1,000 iterations. Even with this reduced sample size, 100% of the iterations retained statistical significance (P < 0.05, median P = 0.0021, interquartile range: 0.001–0.006), confirming that the robust signal in upper glycolysis reflects genuine biological modularity rather than an artifact of statistical power. These results show that diversification of carbon-source utilization exhibits modular patterns that only partially overlap with conventional metabolic classifications.

To ensure that this modular structure was not an artifact driven by global growth rate differences among strains—where inherently fast- or slow-growing isolates could artificially inflate all pairwise correlations—we conducted a sensitivity analysis by controlling for strain-level baselines. Specifically, after carbon-wise standardization (z-score normalization), we subtracted each strain’s mean growth rate across all carbon sources to obtain a normalized growth-rate profile. The interstrain correlation matrix calculated from this normalized profile yielded a dendrogram almost identical to that of the original profile (**Figure S6**). Quantitative topology comparison confirmed extreme structural preservation between the two clustering trees, with an entanglement score of 0.021 (P = 0.001) and a cophenetic correlation coefficient of 0.98 (P = 0.001). These results demonstrate that the observed modularity reflects genuine evolutionary co-variation associated with metabolic architecture rather than strain-level baseline growth differences.

Similarly, the modular structure was not driven by non-growing (zero-growth) entries. Pairs involving zero-growth conditions were extremely rare across all clusters (**Figure S7**). Moreover, re-evaluating hierarchical clustering after excluding zero-growth samples yielded near-identical dendrogram topology (entanglement score = 0.034, P = 0.001; cophenetic correlation = 0.99, P = 0.001; **Figure S7B**–**D**), confirming that the modules arise from quantitative variation in growth rates among growing strains rather than shared non-growth profiles. As a more stringent approach, we also excluded strains containing zero-growth samples on any carbon source. This additional filtration substantially reduced the sample size from 1,732 to 1,325 samples, leaving 53 ECOR strains across 25 carbon sources. The hierarchical clustering based on this strict filtration still exhibited a similar overall dendrogram topology (entanglement score = 0.014, P = 0.001; cophenetic correlation = 0.75, P = 0.001), in which glycolytic and gluconeogenic carbon sources were separated even more clearly (**Figure S7E**–**G**). This analysis confirmed that comparable modules can arise even from this subset of strains.

### Metabolic model captures modular pattern of diversification

To link interstrain correlations to metabolic architecture, we theoretically quantified metabolic dissimilarity among carbon sources. We performed parsimonious flux balance analysis (pFBA) using genome-scale models (GEMs) reconstructed for 70 ECOR strains under standardized carbon-uptake conditions (**Table S5**). We obtained the optimal flux distribution across all reactions for each strain– carbon source pair. To capture internal metabolic architecture, exchange reactions were excluded, and a common set of active internal reactions shared across all models was evaluated (**Table S6**). The flux-based metabolic distance between carbon sources was defined as the cosine distance between their respective flux vectors (**Figure 4A**), and the mean distances across the 70 ECOR models were used for subsequent analyses (**Figure S8**, **Table S7**). A principal coordinate analysis (PCoA) was performed to visualize the overall relationships among carbon sources based on these distances (**Figure 4B**).

**Figure 4.**
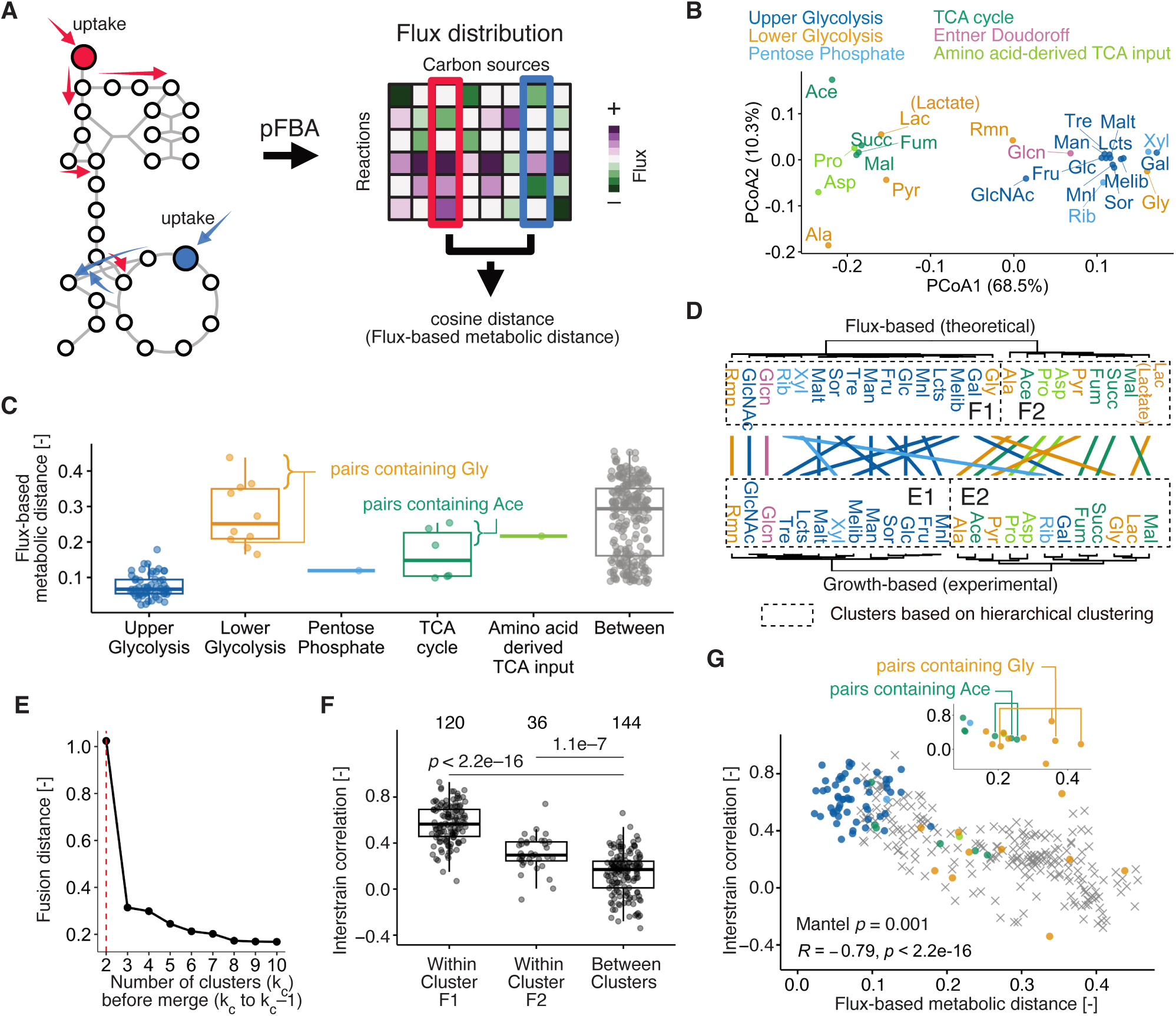
Comparison between theoretical and experimental correlations across carbon sources. (**A**) Schematic representation of metabolic distance calculation. Parsimonious FBA (pFBA) using genome-scale metabolic models identified optimal flux distributions for optimal biomass production. Cosine distance between flux vectors was used as metabolic distance. (**B**) PCoA (Principal Coordinate Analysis) plot based on pairwise mean cosine distances between intracellular flux distributions across all analyzed ECOR strains. Each point represents a distinct carbon source, color-coded according to its metabolic category. The percentage of variance explained by each principal coordinate axis (PCoA1 and PCoA2) is shown in parentheses along the respective axis labels. (**C**) Metabolic distances within conventional pathway categories. Between-category distances are shown as references. (**D**) Tanglegram comparing the flux-based dendrogram (top) and growth-based dendrogram (bottom), both constructed by hierarchical clustering (Ward.D2). (**E**) Scree plot of the dendrogram’s fusion distances across different numbers of clusters (k_c_ prior to merging to k_c_ − 1). The dashed line indicates k_c_ = 2. (**F**) Comparisons of interstrain correlations in ECOR for within- and between-cluster pairs for each cluster. Numbers above the boxes indicate the number of pairs. P values were calculated using the Wilcoxon test (two-sided) with Benjamini-Hochberg (BH) correction. In panels (**C**) and (**F**) boxplot elements follow the same conventions as in Figure 3F. (**G**) Relationship between interstrain correlation and metabolic distance. Colors match pathway categories shown in panel (**C**). Spearman’s R and associated two-sided P-value are indicated. P-value in Mantel test with 999 permutations is indicated. The inset shows pairs after removing upper-glycolysis and between-category pairs.

The flux-based metabolic distances were smaller within upper glycolysis than across categories (**Figure 4C**), whereas TCA-cycle and lower-glycolysis carbon sources showed broad and sometimes large within-category distances. Carbon sources such as glycerol (Gly) and acetate (Ace) were metabolically distant from other members of their assigned categories. These observations indicate that conventional metabolic pathway categories encompass substantial internal heterogeneity, with members of the same category often relying on metabolically distinct reaction sets.

Hierarchical clustering based on these distances closely matched the clustering observed in interstrain correlations (**Figure 4D**). The corresponding tanglegram showed strong concordance between the flux-based and growth-based dendrograms (entanglement score = 0.14, P = 0.022), as did the cophenetic correlation (0.61, P = 0.001). Even after excluding upper-glycolysis carbon sources, the cophenetic correlation remained significant (0.62, P = 0.002), and although the entanglement score improved numerically (0.11 vs 0.14), it did not reach statistical significance (P = 0.089), likely reflecting reduced statistical power with the smaller carbon source set. Nevertheless, the significant cophenetic correlation indicates that overall metabolic architecture, rather than upper-glycolytic topology alone, at least partially contributed to the observed modularity. We also confirmed the relevance of splitting carbon sources into two primary modules̶specifically distinguishing those fueling glycolysis from those feeding gluconeogenesis—as supported by an analysis of the dendrogram’s fusion distances at k_c_ = 2 (**Figure 4E**). These theoretically defined clusters (F1 and F2) were also distinctive from each other in ECOR strains (**Figure 4F**).

Interstrain correlations, as well as PGLS- and PIC-derived evolutionary correlations (**Figure S5**), exhibited a significant negative association with the flux-based metabolic distance (**Figure 4G**; Spearman’s R = −0.79, P < 2.2 × 10^−16^; Mantel test, P = 0.001). This relationship partially accounted for the low correlations observed for within-category pairs such as Ace: although grouped into the same conventional categories, Ace was metabolically distant from others in terms of flux-based metabolic distance (inset of **Figure 4G**). Together, these results demonstrate a close correspondence between patterns of modular diversification in carbon-source utilization and the modular organization of the metabolic network.

As an alternative metric to flux-based metabolic distance, we followed Leiby and Marx^22^ and performed single-reaction knockout simulations to identify essential reactions for optimal growth on each carbon source (**Table S8**). The essentiality-based metabolic distance between carbon sources was defined as the Manhattan distance between their essential-reaction vectors (**Figure S9A**). Median distances across the 70 ECOR models were used. We found that the number of reactions differentiating carbon sources (**Table S9, Figure S9B** and **C**) was small (2–29 reactions depending on carbon source pairs) relative to those essential across all carbon sources (143 reactions), suggesting that this metric tends to reflect differences in catabolic entry reactions for certain pairs. Consistent with this, hierarchical clustering of this metric exhibited no distinctive modules, as indicated by a lack of prominent jumps in the fusion distances (**Figure S9F**). Nevertheless, the essentiality-based metabolic distance was strongly associated with interstrain correlations (Spearman’s R = −0.61, P < 2.2 × 10^−16^; Mantel test, P = 0.001, **Figure S10A**) as well as the overall topological structure of the hierarchical clusters in ECOR strains (entanglement score = 0.11, P = 0.002; cophenetic correlation = 0.59, P = 0.001; **Figure S10C**).

While our theoretical approaches successfully account for the overall modular correlation across carbon sources, it is important to note the current limitations in predicting strain-specific quantitative growth rates directly from GEMs across individual ECOR strains. To directly evaluate model performance, we conducted a pairwise cross-prediction benchmark comparing all strain-specific GEMs against all strain datasets across multiple carbon sources. In this cross-prediction matrix, almost all models consistently demonstrated broad predictive accuracy across carbon sources, confirming that these GEMs successfully capture growth rate differences driven by different carbon sources (**Figure S11**). However, matching model–strain pairs (self-predictions) showed no superior predictive accuracy compared to non-matching pairs, with many models yielding virtually equivalent performance. Furthermore, when controlling for carbon source by examining predictions within individual carbon sources, none of the carbon sources exhibited a statistically significant correlation between predicted and observed growth rates across strains (Spearman’s correlation test, BH-adjusted P ≥ 0.05). Taken together, while GEMs effectively explain growth rate differences across carbon sources, they fail to resolve fine-scale quantitative variations among individual strains.

### Growth-rate profiles in de novo evolution

To test whether modular diversification observed in natural isolates also emerges during de novo evolution, we performed a mutation-accumulation (MA) experiment using a mutator strain (*E. coli* K-12 MG1655*ΔmutS*) propagated solely on glucose (**Figure 5A**). Unlike large-population evolution experiments, MA experiments promote accumulation of spontaneous mutations via genetic drift due to repeated single-colony bottlenecks. If patterns of diversification observed in natural isolates are general, similar covariation in growth rates with metabolic pathway distance should also be observed during de novo evolution.

**Figure 5.**
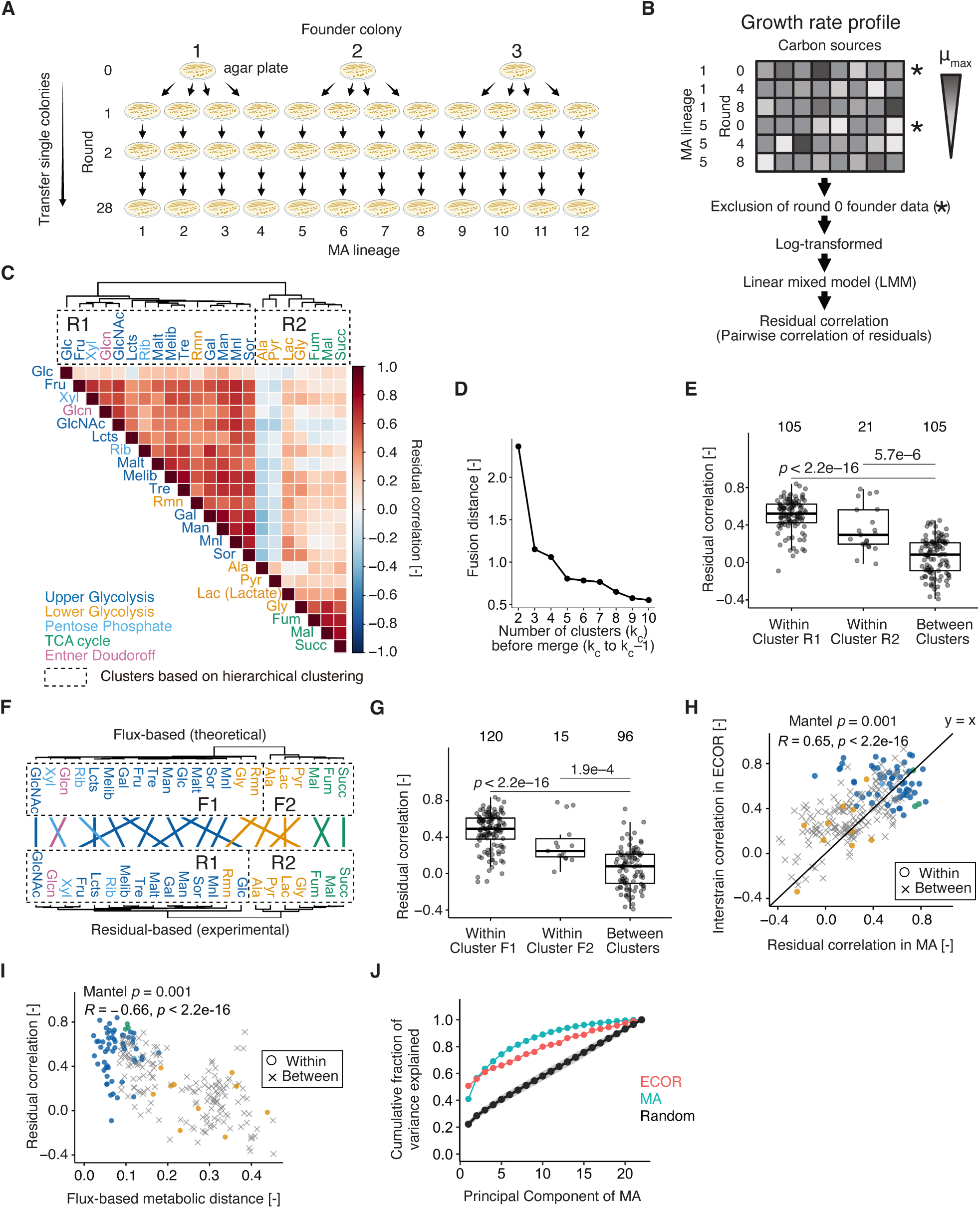
Evolutionary correlations in de novo evolution. (**A**) Schematic of the mutation-accumulation (MA) experiment. Three founders of the mutator strain *E. coli* MG1655 *ΔmutS* were used to initiate MA lineages. A single colony was randomly picked and transferred every two days to fresh M9-glucose agar plates for 28 rounds, generating four MA lineages from each founder. (**B**) Growth-rate profiles for 12 MA lineages across eight time points (0, 4, 8, 12, 16, 20, 24, and 28 rounds). Carbon sources excluded in the ECOR analysis were also excluded here. After removing the founder baseline data (round 0), growth rates were log-transformed and fitted with a linear mixed model (LMM) to account for lineage-specific background and temporal trends. Spearman’s rank correlation coefficients were then computed on the extracted residuals to obtain the residual correlation matrix. (**C**) Heatmap of residual correlations across carbon sources from MA lineages. Dendrogram is based on hierarchical clustering (Ward.D2). Aspartate (Asp), acetate (Ace), and proline (Pro) were excluded because too few MA lineages showed measurable growth. (**D**) Scree plot of the dendrogram’s fusion distances across different numbers of clusters (k_c_ prior to merging to k_c_ − 1). (**E**) Comparisons of residual correlations for within- and between-cluster pairs for each cluster. (**F**) Tanglegram comparing the flux-based dendrogram (top) and residual-based dendrogram (bottom). (**G**) Comparisons of residual correlations for within- and between-cluster pairs for each cluster using the cluster definitions from flux-based metabolic distances (Figure 4D). In panels (**E**) and (**G**) boxplot conventions are identical to those in Figure 3F. Numbers above the boxes indicate the number of pairs. P values were calculated using the Wilcoxon test (two-sided) with Benjamini-Hochberg (BH) correction. (**H**) Relationship between interstrain correlations in ECOR strains and residual correlations in MA lineages. (**I**) Relationship between residual correlation and flux-based metabolic distance in MA lineages. In panels (**H**) and (**I**), Spearman’s R and associated two-sided P-value are indicated. P-value in Mantel test with 999 permutations is indicated. Symbols denote within- (circles) and between-category (crosses) pairs. Colors correspond to pathway categories shown in panel (**C**). (**J**) Cumulative fraction of variance explained when projecting phenotypic variation of ECOR onto the principal component axes derived from MA. The shaded area represents the 95% confidence interval derived from a permutation null model (1,000 iterations) generated by randomly shuffling carbon source values within each ECOR strain.

We quantified OD_max_ and μ_max_ for 12 independent MA lineages across eight time points (rounds 0–28, **Table S4**). We excluded the same carbon sources removed in the ECOR analysis, as well as acetate (Ace), proline (Pro), and aspartate (Asp), because too few MA lineages showed measurable growth (**Figure S12**). Growth-rate profiles were constructed following the same procedure as for the ECOR strains (**Figure 5B**, **Figure S13**). As expected from the design of the MA experiment, μ_max_ varied across both rounds and lineages (**Figure S14**). Loss or gain of metabolic capabilities was rare, consistent with the short timescale. For many carbon sources, variance was lowest in the ancestral state (round 0) and remained elevated thereafter (**Figure S14D**). After removing the founder baseline data (round 0), these growth rates were log-transformed and fitted with a linear mixed model (LMM) to account for founder backgrounds, lineage-specific background, and temporal trends. Log-transformation prevented the LMM from predicting biologically impossible negative growth values. Spearman’s rank correlation coefficients were then computed from the extracted residuals to obtain the residual correlation matrix (hereafter referred to as residual correlations).

### De novo evolution reproduces modular diversification in carbon-source utilization

Hierarchical clustering of residual correlations revealed two major groups (R1 and R2)—one enriched in glycolytic carbon sources and another enriched in TCA-cycle and lower-glycolysis carbon sources (**Figure 5C**), closely resembling the modular structure observed in ECOR strains (**Figure 3D**). The relevance of splitting carbon sources into two primary modules was confirmed by a prominent jump in the dendrogram’s fusion distances at k_c_ = 2 (**Figure 5D**). Within-cluster correlations were significantly higher than between-cluster correlations (**Figure 5E**, Wilcoxon test, BH-adjusted P < 0.05).

Flux-based clustering again captured the experimental clusters in de novo evolution (entanglement score = 0.058, P = 0.001; cophenetic correlation = 0.76, P = 0.001; **Figure 5F**). The clusters (F1 and F2) defined theoretically were also distinctive from each other in de novo evolution (**Figure 5G**). Upper glycolysis carbon sources and intermediates of the TCA cycle exhibited higher residual correlations within each category, while lower glycolysis carbon sources showed lower residual correlations, as observed in ECOR strains (**Figure 5H**). As in ECOR strains, flux-based metabolic distance showed a significant negative association with residual correlations (**Figure 5I**, Spearman’s R = −0.66, P < 2.2 × 10^−16^; Mantel test, P = 0.001). These results indicate that modular diversification patterns observed in natural isolates re-emerge during de novo evolution.

Similarly, the essentiality-based metabolic distance was strongly associated with residual correlations (Spearman’s R = −0.56, P < 2.2 × 10^−16^; Mantel test, P = 0.001, **Figure S10B**) as well as the overall topological structure of the hierarchical clusters in MA lineages (entanglement score = 0.074, P = 0.001; cophenetic correlation = 0.58, P = 0.001; **Figure S10D**), collectively reinforcing the robustness of these findings.

### Limited evidence for historical contingency in lineage-specific evolutionary rates

To assess whether individual lineages followed idiosyncratic evolutionary trajectories — as would be expected under historical contingency — we tested for lineage-specific evolutionary rates using a likelihood ratio test (LRT) comparing random-intercept-only and random-slope LMMs for each carbon source. Of the 22 carbon sources tested, 18 showed no significant lineage divergence in evolutionary rates (LRT, P ≥ 0.05), indicating that parallel evolution predominates across the MA experiment. Significant lineage-specific rates were detected in only 4 carbon sources (Pyr, Xyl, Ala, Lcts), of which only Lcts belongs to the upper glycolytic category; the remainder fall into lower glycolysis and the PP pathway, which require more specialized catabolic routes.

The lineage-specific evolutionary rates (random slopes *b_jk_* in **Equation 1**, **STAR Methods**) showed a significant negative correlation with flux-based metabolic distance across carbon source pairs (Spearman’s R = −0.61, P < 2.2×10^-16^; Mantel test P = 0.001), indicating that lineages evolving faster on one carbon source tended to evolve faster on metabolically similar carbon sources. Thus, even the residual lineage-specific variation reflects metabolic network structure rather than idiosyncratic mutational histories.

To further test whether prior mutations directionally reinforce subsequent evolutionary change, we computed cosine similarities between consecutive within-lineage evolutionary change vectors (**Equation 3**, **STAR Methods**). The observed mean cosine similarity was −0.38 ± 0.37 (SD; n = 60 lineage–step pairs), showing no positive directional alignment and providing no evidence for directional self-reinforcement across successive rounds. Taken together, while lineage-specific evolutionary rates were detectable in a minority of carbon sources (4/22), the predominance of parallel evolution, the metabolic-network-structured pattern of lineage-specific variation, and the absence of directional reinforcement collectively suggest that historical contingency plays a limited role in this MA experiment, and that the metabolic network is the primary detectable organizer of evolutionary trajectories in this experiment.

### Low dimensionality shared between natural and de novo diversifications

Finally, to assess whether patterns identified in de novo diversification are also present in natural variation, we projected ECOR growth-rate profiles onto the principal component (PC) space defined by MA lineages. We then quantified the cumulative fraction of variance in ECOR explained by these MA-derived PCs and compared it with the variance captured within the MA dataset itself, as well as with a permutation null model (**Figure 5J**). The cumulative variance curves for both datasets showed convex upward shapes distinct from random expectations. Notably, the top three PCs of MA lineages explained ∼60% of ECOR variance, demonstrating that both natural and experimental diversification share a low-dimensional structure. Conversely, projecting MA lineages onto the PCA space defined by ECOR strains yielded a comparable result, with the top three ECOR PCs explaining ∼50% of the variance in the MA data. This slight difference may partly reflect the smaller sample size of the ECOR dataset. Taken together, growth rates covary across carbon sources rather than varying independently, revealing modular patterns common to both natural and de novo diversification.

## Discussion

In this study, we asked whether the apparent idiosyncrasy of intraspecific metabolic diversification conceals a recurrent and structured pattern. By integrating quantitative growth phenotyping of natural *E. coli* isolates with de novo laboratory evolution and reaction-level measures of metabolic similarity, we found that diversification in carbon-source utilization is largely independent of phylogeny yet consistently organized into modular and low-dimensional patterns across carbon sources. Notably, closely analogous modular relationships re-emerge during de novo evolution under repeated bottlenecks, and both natural and experimental datasets share a similarly low-dimensional organization of growth-rate variation. Together, these observations show that metabolic phenotypes vary along a limited set of recurring, module-linked axes that are reproducible across evolutionary contexts.

Why, then, did earlier studies not report modular patterns like those observed here? A key reason is that previous works were not designed to capture quantitative covariation in growth rates across a broad range of carbon sources. For instance, Sabarly et al.^20^ relied on Biolog assays that measure cellular respiration rather than growth rate itself, and respiration signals are known to be an unreliable quantitative proxy for growth rates ^22^, especially within species. Likewise, the LTEE dataset analyzed by Leiby and Marx^22^ was optimized to study long-term adaptation to glucose, and the ancestral LTEE strains lacked growth on many TCA-cycle carbon sources, effectively removing an entire metabolic module from phenotypic space. As a result, these results did not provide the combination of quantitative growth-rate measurements and metabolic breadth needed to reveal modular covariation across carbon sources. By contrast, our study combines high-resolution growth measurements with evolutionary variation across both natural isolates and de novo-evolved lineages, while retaining a wide range of metabolic capabilities. This combination of quantitative resolution and metabolic breadth makes previously latent structure visible, clarifying how modular patterns of diversification can be observed within species.

One possible explanation for the recurrent modular diversification observed here is the accumulation of mutations directly affecting reactions specific to individual metabolic modules. Under this view, repeated perturbation of the same reactions or pathways could generate coordinated changes in growth performance across subsets of carbon sources. However, several considerations suggest that this mechanism alone is unlikely to fully explain the observed patterns. The number of reactions differentiating carbon sources is small (approximately 30 reactions, **Figure S9B**–**D**) relative to genome size (4,000–5,800 genes), and many essential reactions are shared broadly across carbon sources. Moreover, the number of substitutions accumulated over the timescale of the mutation-accumulation experiment is limited (approximately 30 substitutions per genome over 30 rounds estimated previously^31^), making repeated direct hits on module-specific components across independent lineages improbable. While direct effects on individual reactions may contribute in certain cases, they are therefore unlikely to account for the recurrent and parallel diversification patterns observed across evolutionary contexts.

A complementary and hypothetical scenario is that mutations influence growth phenotypes indirectly through higher-level regulatory or network-mediated processes. In *E. coli*, a small number of global transcriptional regulators respond sensitively to diverse physiological perturbations^5^ including genetic mutations, and can propagate their effects to diverse sets of genes across the genome, including metabolic genes involved in reactions belonging to different metabolic modules. Under this hypothesis, modular diversification need not arise from modular transcriptional regulation per se. Instead, perturbations introduced by a small number of genetic changes may propagate system-wide through transcriptional regulatory networks, and their downstream phenotypic effects become organized into modular patterns by the structure of the metabolic network. Changes affecting such regulatory layers could therefore shift flux distributions across groups of reactions, even in the absence of coordinated gene-expression changes aligned with metabolic pathways. From this perspective, distinct genetic changes need not repeatedly affect the same reactions to generate similar growth-rate correlations across carbon sources. Instead, different mutations may lead to coherent, module-level shifts in metabolic activity that reflect the structure of the underlying network. This interpretation is consistent with the observed alignment between modular diversification patterns and reaction-level measures of metabolic similarity, without requiring a direct one-to-one correspondence between specific mutations and individual reactions.

We emphasize that this mechanistic scenario remains a working hypothesis, as confirming the exact mutational targets will require full-genome sequencing of the MA lineages in future studies. Nevertheless, this network-level view also provides a unified interpretation of the two defining features of our data: the parallel re-emergence of similar diversification patterns across evolutionary contexts and the weak correspondence between metabolic phenotypes and phylogeny. The recurrence of modular and low-dimensional structures in both natural isolates and de novo-evolved lineages does not require convergent selection acting on the same carbon sources or identical genetic changes. Rather, it may reflect repeated exploration of a shared phenotypic space structured by metabolic interactions. When phenotypic variation is concentrated along a limited number of network-defined axes, distinct mutations and evolutionary histories can readily give rise to similar patterns of covariation across growth conditions. In such a low-dimensional space, phylogenetically distant strains may occupy nearby phenotypic regions, while closely related strains may diverge along different axes. From this perspective, weak phylogenetic signal and structured, recurrent diversification are not contradictory but mutually compatible outcomes of metabolic organization shaping the space of accessible phenotypic variation.

Our findings also provide a critical link to recent macro-evolutionary studies on bacterial carbon preference while offering a fundamentally distinct perspective. Recent large-scale phenotyping across diverse heterotrophic bacteria demonstrated that metabolic preferences organize along a few major axes—predominantly separating sugar and organic-acid utilization—and can be predicted from gross genomic features such as pathway gene counts and GC content ^23^. Because these broad genomic features vary minimally within a single species, such models cannot account for the structured quantitative variation we observe among closely related *E. coli* strains. Strikingly, our results show that a closely analogous modular split between glycolytic substrates and organic acids re-emerges both across natural *E. coli* isolates and during de novo laboratory evolution. This demonstrates that low-dimensional, module-linked diversification does not strictly depend on large-scale variations in gene content or GC content. This convergence with macro-evolutionary patterns is consistent with the finding that metabolic distances derived from genome-scale modeling capture ecologically relevant structure at the family level among Proteobacteria ^24^, and extends that principle to within-species metabolic diversification: the shared architecture of the metabolic network itself exerts a strong constraint, guiding quantitative phenotypic variation along recurring axes even within the tight genomic boundaries of a single species.

In conclusion, our results show that intraspecific diversification of carbon-source utilization in *E. coli* is neither random nor strongly constrained by phylogeny, but instead repeatedly organizes into modular and low-dimensional patterns that re-emerge across natural and experimental evolutionary contexts. By combining quantitative growth phenotyping with network-level measures of metabolic similarity, we reveal an organizational structure underlying metabolic variation that is not apparent from gene content or conventional pathway classifications alone.

At the same time, our analyses are restricted to a single species and to growth-rate phenotypes across defined carbon sources, and they do not identify the specific molecular mechanisms generating these patterns. Dissecting how genetic changes map onto the observed module-level structures— through regulatory interactions, metabolic flux reorganization, or other processes—remains an important goal for future work. More broadly, this study illustrates how integrating quantitative phenotyping with network-based perspectives can uncover structured patterns of variation within species, even when evolutionary change appears weakly coupled to phylogeny.

Several factors likely account for the inability of the constructed ECOR GEMs to predict strain-specific quantitative growth rates. First, automatically generated draft models lack the extensive, manual curation that defines highly benchmarked models like iML1515, a limitation compounded by the fragmented, multi-contig nature of available draft genome assemblies for the ECOR collection. Tuning these models to capture subtle quantitative phenotypes would require fitting them to the very growth data collected here, which would introduce circular reasoning. Second, standard flux balance analysis assumes optimal flux allocation based solely on gene presence–absence, inherently omitting regulatory constraints such as variations in gene expression levels or non-synonymous mutations altering enzyme kinetics. Indeed, our previous work showed that expression levels of genes involved in central metabolism—including amino acid metabolism and the TCA cycle—vary substantially across strains despite high gene conservation ^5^. Integrating quantitative strain-specific expression profiles or multi-omics data with GEMs will be essential in future studies to fully bridge the gap between genotype and quantitative metabolic phenotypes at the intraspecific level.

## Resource availability

### Lead contact

Requests for further information, resources, and reagents should be directed to and will be fulfilled by the lead contact, Chikara Furusawa.

### Materials availability

All strains generated in this study are available upon request from the lead contact. Note that distribution of certain strains may be subject to third-party Material Transfer Agreement (MTA) restrictions.

### Data and code availability

The raw OD data and GEMs are available at Zenodo (https://doi.org/10.5281/zenodo.19491366). Custom scripts used in this study are available at GitHub (https://github.com/tsuruubi/ECORMA).

## Acknowledgements

We thank H. Koike for technical support in MA experiments. We thank Shannon Manning at Michigan State University for providing the ECOR strains. This work was supported by RIKEN Junior Research Associate Program (to K.S.), Japan Society for the Promotion of Science (JSPS) KAKENHI (18H02427, 24K21985 to S.T.; 17H06389 to C.F. and S.T.; 22K21344, 23K27164, and 24H01798 to C.F.), Japan Science and Technology Agency (JST) ERATO (JPMJER1902 to S.T. and C.F.).

## Author contributions

Conceptualization, S.T.; Methodology, S.T. and C.F.; Investigation, K.S. and S.T.; Data Curation, K.S. and S.T.; Formal Analysis, K.S. and S.T.; Validation, S.T.; Visualization, K.S. and S.T.; Resources, S.T. and C.F.; Software, K.S. and S.T.; Funding Acquisition, S.T. and C.F.; Project Administration, S.T.; Supervision, S.T. and C.F.; Writing – Original Draft, K.S.; Writing – Review & Editing, K.S., S.T., and C.F.

## Declaration of generative AI and AI-assisted technologies in the writing process

During the preparation of this manuscript, the authors used Claude Sonnet 4.6 (Anthropic) to polish the manuscript for clarity, coherence, and conciseness. After using this tool, the authors reviewed and edited the content as needed and take full responsibility for the content of the publication.

## Supplemental information

Document S1. Figures S1–S14

Spreadsheet S1. Tables S1–S9

## STAR Methods

### Key resources table

| REAGENT or RESOURCE | SOURCE | IDENTIFIER |
|---|---|---|
| Bacterial and virus strains |  |  |
| *E. coli* reference (ECOR) collection | Ochman and Selander<sup>25</sup> | ECOR01–72; |
| *E. coli* K-12 MG1655 *ΔmutS* | This study | N/A |
| Chemicals, peptides, and recombinant proteins |  |  |
| Bulk chemical reagents for M9 minimal medium and carbon sources | This study / Various | See Supplementary Table S2 |
| LB Broth, Miller | BD Biosciences | Cat# 244620 |
| Agar, Powder | Fujifilm Wako | Cat# 010-08725 |
| Chloramphenicol | Fujifilm Wako | Cat# 034-10572 |
| L (+)-Arabinose | Fujifilm Wako | Cat# 012-04581 |
| Ampicillin Sodium | Fujifilm Wako | Cat# 012-23303 |
| Deposited data |  |  |
| Raw OD data | This study | https://doi.org/10.5281/zenodo.19491366 |
| Genome-scale metabolic model of 70 ECOR strains | This study | https://doi.org/10.5281/zenodo.19491366 |
| Recombinant DNA |  |  |
| pKD46 | Datsenko & Wanner<sup>32</sup> | N/A |
| pKD32 | Datsenko & Wanner<sup>32</sup> | N/A |
| Software and algorithms |  |  |
| Custom Python scripts | This study | https://github.com/tsuruubi/ECORMA |
| Python v3.9.20 | Python Software Foundation | https://www.python.org/; RRID:SCR_008394 |
| COBRApy v0.29.0 | Ebrahim et al. <sup>33</sup> | https://github.com/opencobra/cobrapy; RRID:SCR_019500 |
| NumPy v2.0.2 | Harris et al. <sup>34</sup>; NumPy Development Team | https://numpy.org/; RRID:SCR_008633 |
| Pandas v2.2.3 | Pandas Development Team | https://pandas.pydata.org/; RRID:SCR_018214 |
| SciPy v1.13.1 | Virtanen et al. <sup>35</sup>; SciPy Development Team | https://scipy.org/; RRID:SCR_008058 |
| IBM ILOG CPLEX Optimization Studio | IBM Corp. | https://www.ibm.com/products/ilog-cplex-optimization-studio |
| CarveMe | Machado et al. <sup>36</sup> | https://github.com/cdanielmachado/carveme; RRID:SCR_028775 |
| Custom R scripts | This study | https://github.com/tsuruubi/ECORMA |
| R, v4.5.2 | R Core Team<sup>37</sup> | https://www.r-project.org/; RRID:SCR_001905 |
| growthcurver, v0.3.1 | Sprouffske & Wagner<sup>38</sup> | https://github.com/sprouffske/growthcurver |
| ape, v5.8-1 | Paradis & Schliep<sup>39</sup> | https://cran.r-project.org/package=ape; RRID:SCR_002092 |
| caper, v1.0.4 | Orme et al. <sup>40</sup> | https://cran.r-project.org/package=caper; RRID:SCR_027575 |
| phytools, v2.5-2 | Revell<sup>41</sup> | https://github.com/liamrevell/phytools; RRID:SCR_017249 |
| dendextend, v1.19.1 | Galili<sup>42</sup> | https://github.com/talgalili/dendextend/; RRID:SCR_026116 |
| vegan, 2.7-5 | Oksanen et al. <sup>43</sup> | https://cran.r-project.org/package=vegan; RRID:SCR_011950 |
| lme4, v2.0-1 | Bates et al. <sup>44</sup> | https://cran.r-project.org/package=lme4; RRID:SCR_015654 |
| Other |  |  |
| Biomek NXp span-8 laboratory automation workstation | Beckman Coulter | Biomek NXp span-8, Cat# A31840; RRID:SCR_019625 |
| STX44 shaking incubator | Liconic | STX44, Cat# 9118-09-00 |
| FilterMax F5 multi-mode microplate reader | Molecular Devices | FilterMax F5, Cat# |
| LPX220 microplate hotel | Liconic | LPX220, Cat# 9144-00-50 |
| Teleshake Magnetic Shaking System | Thermo Scientific | Cat# 50108900 |
| DWMax M-BR-034P microplate shaker | Taitec | MBR-034P |
| Infinite F200 multimode plate reader | Tecan | Infinite F200 |
| Nunc MicroWell 96-well Nunclon Delta-Treated, Flat-Bottom Microplate | Thermo Scientific | Cat# 167008 |
| Corning 384-well Clear Flat Bottom Polystyrene Not Treated Microplate | Corning | Cat# 3680 |

### Experimental model and study participant details

#### Bacterial strains

We used a collection of 72 natural isolates of *Escherichia coli* from the ECOR (*E. coli* reference) collection^25^. These strains were originally isolated from diverse hosts and geographic regions and cover the seven major *E. coli* phylogroups (**Table S1**) ^26^. *E. coli* K-12 MG1655 *ΔmutS* was constructed by standardized λ -red homologous recombination using the pKD46 plasmid ^32^ in the presence of 60 mM arabinose (FUJIFILM Wako, Osaka, Japan, Cat. No. 012-04581) and 50 μg/mL ampicillin (FUJIFILM Wako, Osaka, Japan, Cat. No. 012-23303), as detailed previously ^5^. The chloramphenicol resistance gene on the pKD32 plasmid ^32^ was amplified by PCR with the same chimeric primers as those used to knock out *mutS* in a previous study^45^ and used as a selection marker for *mutS* deletion in the presence of 30 μg/mL chloramphenicol (FUJIFILM Wako, Osaka, Japan, Cat. No. 034-10572). The resultant strain was used as the ancestral strain for the mutation-accumulation (MA) experiment. Deletion of the *mutS* gene, a key component of the mismatch-repair system, substantially elevates the mutation rate relative to wild type^46^, and previous work estimated a mutation rate of approximately 10 ^−8^ substitutions per base pair per generation^31^.

### Method details

#### Culture media and carbon sources

All reagents and suppliers used in this study are listed in **Table S2**. Growth assays were conducted in a synthetic medium derived from M9 minimal medium (pH 7.2), detailed previously^47^. In short, the synthetic medium contains 47.7 mM Na_2_HPO_4_, 22.0 mM KH_2_PO_4_, 8.6 mM NaCl, 18.7 mM NH_4_Cl, 0.5 mM MgSO_4_, 0.1 mM CaCl_2_, 10 μM FeSO_4_, trace elements (3.4 nM (NH_4_)_6_Mo_7_O_24_, 466 nM H_3_BO_3_, 64 nM CoCl_2_, 17 nM CuSO_4_, 94 nM MnCl_2_, 11 nM ZnSO_4_), and 10 μM vitamins (thiamine, pantothenate, p-aminobenzoic acid, p-hydroxybenzoic acid, 2, 3-dihydroxybenzoic acid). Each carbon source was supplied individually at a final concentration of 5 mM. In the MA experiment, the synthetic medium was solidified with 1.5% agar (FUJIFILM Wako, Osaka, Japan, Cat. No. 010-08725), and glucose was provided as the sole carbon source at 25 mM. Carbon sources were assigned to conventional metabolic pathway categories^30^, including upper glycolysis (glucose, galactose, mannose, fructose, maltose, trehalose, lactose, melibiose, sorbitol, mannitol, and n-acetyl-d-glucosamine), lower glycolysis (rhamnose, glycerol, lactate, pyruvate, serine, threonine, alanine), the TCA cycle (acetate, fumarate, malate, succinate, and citrate), the Entner–Doudoroff (ED) pathway (gluconate), and the pentose phosphate (PP) pathway (ribose and xylose). Amino acids that feed into the TCA cycle without passing through these central pathways were grouped together as amino-acid–derived TCA inputs (proline, aspartate, glutamate, arginine, histidine, and lysine).

#### MA experiment

The MA experiment was conducted following the procedures detailed previously^5^. In brief, three independent colonies of MG1655 *ΔmutS*, designated founders 1, 2, and 3, were used to initiate lineages, thereby minimizing the influence of pre-existing mutations. Four independent lineages were generated from each founder, yielding twelve lineages in total. Each lineage was propagated through repeated single-colony bottlenecks: every two days, a randomly chosen colony was transferred to a fresh agar plate. Plates were incubated at 32°C until the next transfer, and this process was repeated for 28 rounds. Samples from rounds 0, 4, 8, 12, 16, 20, 24, and 28 were preserved at −80°C. Approximately one base-pair substitution is expected to become fixed per round in this system, based on estimates from a previous MA experiment^31^.

#### Growth assay design

Growth assays for the ECOR strains and MA lineages were conducted inside a clean booth using an automated culture system (Biomek NX Span-8; Beckman Coulter, Brea, CA, USA) integrated with a plate reader (FilterMax F5; Molecular Devices, San Jose, CA, USA), a plate shaker (StoreX STX44; Liconic, Mauren, Liechtenstein), a plate hotel (LiCotel LPX220; Liconic), and a Teleshake module (Thermo scientific, Waltham, MA, USA, Cat. No. 50108900)^27^. The incubation temperature was maintained at 33°C to ensure stable operation and minimize evaporation^48^. Frozen glycerol stocks were revived by inoculation into lysogeny broth (LB; BD Biosciences, Franklin Lakes, NJ, USA, Cat. No. 244620) in a 96-well microplate (Thermo scientific, Cat. No. 167008) and incubated in an incubator (DWMax M-BR-034P; Taitec, Saitama, Japan) at 32°C for approximately 24 hours with shaking at 800 rpm. From each preculture, after measuring OD using a plate reader (Infinite F200; Tecan, Switzerland), 4 μL of culture was added to 800 μL of sterile M9 buffer to generate diluted inocula. These inocula (15 μL/well) were mixed with the synthetic medium (65 μL/well) containing a single carbon source (resulting in a 1,074-fold dilution), and four biological replicates were prepared per strain–carbon combination. Each culture was established in a 384-well microplate (Corning Inc., NY, USA, Cat. No. 3680) with a total volume of 80 μL/well and a path length of 8.7 mm (**Table S3**). Plates were loaded into the automated platform and incubated without shaking, and OD₆₀₀ was measured every hour for 72 hours. To avoid the influence of cell settling prior to each measurement, plates were briefly agitated for one minute.

### Quantification and statistical analysis

#### Growth curve fitting and classification

Growth curves were analyzed by R^37^ using the growthcurver package^38^. OD measurements between one and three hours were excluded because of their high variability due to instrumental initial instability. Cultures with an initial OD ≥ 0.07 were excluded owing to inoculation errors or the presence of bubbles. The remaining measurements were fit to a logistic growth model using the SummarizeGrowth function, yielding estimates of the maximum growth rate (μ_max_), carrying capacity (K), and the coefficient of determination (R²). To avoid contaminating the fits with measurements from the death phase, we calculated the moving-median OD and identified its maximum; data points collected after the time at which OD first dropped below 90% of this maximum were excluded from curve fitting. We then evaluated the maximum observed OD (OD_max_) relative to K to classify growth outcomes for each culture (**Figure S1** and **S2**). Cultures with OD_max_ ≤ 0.1 were classified as exhibiting “No growth”, and for these cultures OD_max_ was reset to zero. When OD_max_ was lower than 80% of K, the culture was considered to show “Slow/Partial growth”. Cultures that reached at least 80% of K were considered to show “Growth”. This three-category classification served solely as a data quality indicator; the distinction between Growth and Slow/Partial growth did not affect downstream analyses, as μ_max_ was used as a continuous variable for all cultures in either category.

Each strain–carbon combination was represented by four replicative cultures. If three or more replicates fell into the same growth category, that category was assigned as the representative category for that strain–carbon combination. When no such majority existed, the combination was assigned to the “Unstable” category. This label serves solely as a data quality indicator and did not affect downstream calculations. Representative OD_max_ and μ_max_ values were calculated as the mean of replicates not classified as “No growth,” regardless of whether the combination had a majority category or was labeled “Unstable” (**Figure S3A**). For combinations whose representative category was “No growth,” both representative OD_max_ and μ_max_ were set to zero. All downstream analyses used μ_max_ from these representative values (**Table S4**). Two strains (ECOR29 and ECOR52) exhibited little or no growth on over half of the carbon sources, likely due to auxotrophy, and were excluded from downstream analyses. Some carbon sources showed little or no growth across most ECOR strains, especially amino acids such as lysine and histidine (**Figure S2**). To ensure sufficient data coverage, we retained only carbon sources that supported Growth or Slow/Partial growth in at least 80% of the 70 remaining ECOR strains. For strain–carbon combinations classified as Growth or Slow/Partial growth, the corresponding μ_max_ value was used; μ_max_ was set to zero for No-growth combinations. This procedure yielded growth-rate profiles for 70 ECOR strains across 25 carbon sources (**Table S4**).

To quantify the similarity in growth-rate profiles between strain pairs, we calculated Spearman’s rank correlation of μ_max_ across carbon sources for each pair of ECOR strains; this pairwise correlation is referred to as growth similarity throughout. Separately, to quantify the degree to which growth rates on two carbon sources co-vary across strains, we calculated Spearman’s rank correlation of μ_max_ across the 70 ECOR strains for each pair of carbon sources; this pairwise correlation is referred to as the interstrain correlation throughout.

#### Phylogenetic reconstruction of ECOR strains

Maximum-likelihood phylogenetic tree of the ECOR collection was retrieved from a previous study^26^. This tree was inferred from core-genome alignments (758,341 bp) of 125 *E. coli* strains, including all ECOR isolates and K12-MG1655 strain analyzed here. Core-genome SNPs identified using Parsnp^49^ served as input for PhyML^50^ under the HKY85 model. Phylogroup (A, B1, B2, C, D, E, and F) assignments followed those of the published tree. Pairwise patristic distances were calculated from the phylogenetic tree and used as measures of phylogenetic distance in downstream analyses of metabolic diversification. The association between phylogenetic distance and growth similarity was assessed using the Mantel ^28^ test (999 permutations) implemented in the R package vegan ^43^, with Spearman’s rank correlation reported as the effect size. Phylogenetic signal (Blomberg’s K^29^) was estimated using the phylosig function in the R package phytools^41^. Branches with lengths below 1×10⁻⁶ were adjusted to this minimum value to prevent numerical instability. Blomberg’s K value of 1 indicates that a trait’s evolutionary similarity across species perfectly matches what is expected under a random Brownian motion model of evolution.

#### Accounting for phylogenetic structure and time-series dependence in MA lineages

To account for the non-independence of observations arising from repeated measurements across time points (rounds 4–28) and the nested evolutionary structure (lineages derived from distinct founders), we analyzed log-transformed growth rates of the MA experiment using linear mixed-effects models (LMMs) implemented in the lme4 ^44^ R package. Log-transformation was applied to prevent the model from predicting biologically impossible negative growth values—which can occur under standard linear models when performance declines over evolutionary time—thereby constraining fitted trajectories to biologically plausible positive values. Specifically, for each carbon source *k*, the log-transformed growth rate *Y_ijkt_* of lineage *j* (*j* ∈ {1,2,⋯,12}) derived from founder *i* ( *i* ∈ {founder 1, founder 2, founder 3}) at round *t* (*t* ∈ {4,8,⋯,28}) was modeled as:

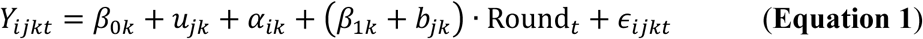

where *β_0k_* is the overall fixed intercept, *α_ik_* captures the fixed effect of founder background *i*, and 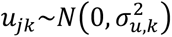 accounts for the lineage-specific random intercept following a normal distribution *N* with mean zero and variance 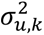. To avoid overfitting or singular fits due to the small number of founders (N = 3), founder background was included as a fixed effect rather than a nested random effect, providing equivalent baseline control. *β_1k_* is the overall fixed slope for evolutionary time (Round*_t_*), while 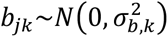 captures the lineage-specific random slope. The combined value *β*_1*k*_ + *b*_j*k*_ defines the net evolutionary rate for lineage *j* on carbon source *k*. 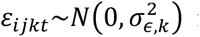 represents the residual error, capturing condition-specific phenotypic fluctuations after removing all baseline, lineage-level, and temporal trends. In cases where random-slope models exhibited singular fits or non-convergence, a fixed-effects interaction model of the same structure — treating *u*_j*k*_ and *b*_j*k*_ as fixed parameters — was fitted as a backup via standard linear regression. Using the resulting residual matrix (samples×carbon sources), we recalculated pairwise Spearman’s rank correlations across carbon sources to obtain residual correlation and tested their association with flux-based metabolic distance using the Mantel test (999 permutations), with Spearman’s rank correlation reported as the effect size.

#### Testing for lineage-specific evolutionary rates and historical contingency

To test whether individual lineages exhibited idiosyncratic evolutionary rates — as would be expected under historical contingency — we performed likelihood ratio tests (LRTs) for each carbon source by comparing the full random-slope model (**Equation 1**) against a reduced random-intercept-only model:

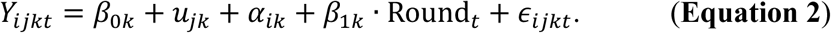

Both models were fitted by maximum likelihood (REML = FALSE) to enable valid likelihood ratio comparisons. A significant LRT (p < 0.05) was taken as evidence that the variance of lineage-specific slopes 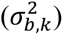 is significantly greater than zero, indicating divergent evolutionary rates across lineages. For carbon sources where the random-slope model returned a singular fit, the LRT was not applicable; in these cases, the random-slope variance converged to zero 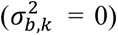, which was interpreted as evidence of negligible lineage divergence.

To examine whether the pattern of lineage-specific evolutionary rates is structured by the metabolic network, we computed pairwise Spearman’s rank correlations among the net evolutionary rates (*β*_1*k*_ + *b*_j*k*_) across all carbon source pairs and tested their association with flux-based metabolic distance using the Mantel test (999 permutations), with Spearman’s rank correlation reported as the effect size.

To further test whether prior mutations directionally reinforce subsequent evolutionary change within individual lineages, we computed the cosine similarity between consecutive within-lineage evolutionary change vectors. For each lineage *j* and each consecutive round pair (*t*, *t* + 4), the change vector was defined as Δ*ε*_j*kt*_ = *ε_i_*_j*kt*_ − *ε_i_*_j*k*(*t*-4)_ across all carbon sources *k*. Cosine similarity between vectors at steps (*t* − 4, *t*) and (*t*, *t* + 4) was calculated as:

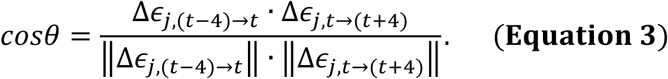

The mean and standard deviation of cosine similarities were computed across all lineage–step pairs (n = 60) to summarize the distribution of observed directional change.

#### Phylogenetically controlled evolutionary correlations in ECOR strains

To account for potential phylogenetic confounding when evaluating evolutionary correlations across carbon sources in ECOR collection, we calculated phylogenetic signal-adjusted correlations using two complementary approaches: Phylogenetic Generalized Least Squares (PGLS ^51^) and Phylogenetically Independent Contrasts (PIC ^52^). PGLS assumes normality of residuals, whereas PIC-based Spearman correlation provides a non-parametric robustness check against this assumption. Prior to these analyses, growth rate profile matrix (*μ_max_* across strains and carbon sources) was column-wise standardized (z-score normalized) for each carbon source.

In PGLS analysis, relative growth traits across strains were fitted using Generalized Least Squares (pgls function in the caper ^40^ R package) for each pair of carbon sources. Pagel’s *λ* was estimated via maximum likelihood (ML) directly from the data alongside the regression parameters (lambda = “ML”), allowing the degree of phylogenetic correction to be adaptively determined. For certain carbon source pairs where ML optimization failed to converge due to highly depleted phylogenetic signals, *λ* was fixed at a negligible value (10 ^−6^), rendering the model effectively equivalent to a standard ordinary least squares regression. For each carbon source pair, the adjusted correlation coefficient *r* was derived from the *t*-statistic of the slope parameter (*t_stat_*) as:

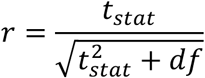

where *df = n-2*, with *n* being the number of strains with valid growth measurements for both carbon sources.

As an alternative non-parametric phylogenetic control, independent contrasts for each carbon source were calculated along the maximum likelihood phylogenetic tree of the ECOR strains using Felsenstein’s method (pic function in the ape ^39^ R package). Pairwise evolutionary correlations between carbon sources were then evaluated using Spearman’s rank correlation coefficients on the resulting contrast matrix across all valid traits.

#### Statistical comparison of metabolic distances across pathway categories

For each of the six conventional metabolic pathway categories, pairwise interstrain correlations were classified as intra-category (both carbon sources belonging to the same category) or inter-category (carbon sources from different categories). Within each category, intra-versus inter-category correlations were compared using a Wilcoxon rank-sum test, with P-values adjusted for multiple comparisons using the Benjamini-Hochberg (BH) procedure (**Figure 3G**). To confirm that the significance observed for upper glycolysis was not an artifact of its larger sample size relative to other categories, four carbon sources were randomly drawn from upper glycolysis (matching the number of TCA-cycle substrates) over 1,000 iterations and the same Wilcoxon test was applied in each iteration.

#### Construction of genome-scale metabolic models

Genome-scale metabolic models were constructed for all 70 ECOR strains (**Table S1**). Protein sequences of these strains were obtained from a previous study^53^ and used as input for CarveMe ^36^ to generate draft models. The models were then gap-filled on in silico LB medium and M9 medium containing glucose (deposited in Zenodo). The genome-scale metabolic model iML1515 for K12-MG1655 strain was obtained from a previous study ^54^.

#### Estimation of metabolic distance

Flux balance analysis (FBA) was conducted using the genome-scale metabolic models implemented in COBRApy^33^. All exchange reactions were initially closed. Subsequently, the oxygen exchange reaction was opened with a lower bound of −10 mmol gDCW⁻¹ h⁻¹, whereas those for inorganic ions, vitamins, and trace metals were opened with a lower bound of −1000 mmol gDCW⁻¹ h⁻¹ to emulate M9 minimal medium (**Table S5**). To standardize carbon availability, we used a carbon-number-normalized uptake scheme^22^, setting the lower bound of each carbon-source exchange reaction to−(60/nC), where nC is the number of carbon atoms in the carbon source. This scheme fixed total available carbon at 60 mmol C gDCW⁻¹ h⁻¹ for all carbon sources, which corresponds to 10 mmol gDCW⁻¹ h⁻¹ in the case of glucose. The biomass reaction (the Growth reaction for the models constructed by CarveMe; BIOMASS_Ec_iML1515_WT_75p37M for iML1515) served as the optimization objective. The maximal biomass production rate was computed by FBA using the CPLEX solver (IBM) and denoted as Z*. Of the carbon sources tested, 30 supported positive biomass production; histidine and lysine did not and were omitted from further analyses.

To evaluate metabolic distance between carbon sources based on predicted intracellular flux distributions, we performed parsimonious FBA (pFBA ^55^) using the genome-scale metabolic models of all analyzed ECOR strains and the iML1515 model. pFBA was executed under the same growth medium constraints and carbon-number-normalized uptake rate as described above. For each strain–carbon source pair, we obtained the optimal flux distribution across all reactions. To compare flux distributions across strains and conditions on a consistent basis, exchange reactions (EX_) were excluded, and a common set of active internal reactions shared across all models was identified. Reactions that maintained zero flux across all evaluated conditions were removed, resulting in a trimmed flux matrix (**Table S6**). For each individual strain, pairwise similarity between flux distributions corresponding to different carbon sources was calculated using cosine similarity:

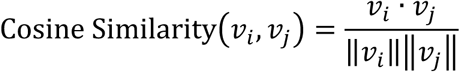

where *v_i_* and *v_j_* represent the intracellular flux vectors for carbon sources *I* and *j*, respectively. The flux-based metabolic distance was then defined as the cosine distance (1−Cosine Similarity). Finally, the mean cosine distances across all analyzed ECOR strains were computed for each carbon-source pair and used for downstream comparisons (**Table S7**). Principal coordinate analysis (PCoA) was applied to the mean cosine distance matrix using the cmdscale function in R.

To determine essential reactions for optimal growth (Z*) under each condition, we performed in silico single-reaction knockouts by setting both bounds of each reaction to zero and re-optimizing FBA under the same medium composition. If biomass production dropped below 0.999 of its optimal value (i.e. 0.999 × Z*), the reaction was deemed essential; otherwise, it was classified as non-essential. This yielded a binary matrix of essential reactions across carbon sources (**Table S8**). The essentiality-based metabolic distance between any two carbon sources was defined as the Manhattan distance between their essential-reaction vectors. The reactions that contribute to distances between carbon sources were listed in **Table S9** for each strain. The median distance across all 70 ECOR models was used for further analyses (**Table S7**).

The associations between metabolic distances (flux-based and essentiality-based) and growth-based correlations (interstrain correlation, PIC- and PGLS-derived evolutionary correlations for the ECOR dataset; residual correlations and lineage-specific slope correlations for the MA dataset) were assessed using the Mantel test (999 permutations), with Spearman’s rank correlation reported as the effect size.

#### Cross-prediction benchmark of strain-specific GEMs

To evaluate the strain-specificity of GEM-based growth predictions, we constructed a cross-prediction matrix by computing Spearman’s rank correlation between FBA-predicted optimal biomass production flux (Z*) and experimentally measured μ_max_ for all combinations of ECOR GEMs and ECOR strain datasets across carbon sources. For the iML1515 reference model, the mean μ_max_ of the MA ancestor (round 0) was used as the experimental reference. Self-prediction performance (matching model–strain pairs) was compared with cross-strain prediction performance (non-matching pairs) by computing a specificity index defined as the difference between self-prediction Spearman’s R and the mean cross-strain Spearman’s R. Additionally, within-carbon-source Spearman’s rank correlations between predicted and measured growth rates across strains were computed, with P-values adjusted using the BH procedure (**Figure S11**).

#### Construction and comparison of dendrograms

For each pair of strains or carbon sources, we computed Spearman’s rank correlation coefficient (R) for μ_max_ values. Distances were defined as 1 − Spearman’s R, and hierarchical clustering was performed using the hclust function in R with Ward.D2 method. A similar procedure was applied to metabolic distances derived theoretically. To determine the number of distinct clusters (modules, k_c_), we analyzed the fusion distances obtained from hierarchical clustering across a range of cluster numbers (k_c_ = 2 to 10). The number of clusters was determined to be k_c_ = 2 based on a prominent increase in fusion height, indicating that merging the two main clusters would incur a substantial loss of distinct cluster structure.

To compare dendrograms, we generated tanglegrams using the R package dendextend ^42^. Trees were first untangled using the untangle function with the “step2side” method to minimize line crossings. Entanglement scores were calculated for these untangled trees, and significance was assessed using permutation tests in which tip labels were randomly shuffled 1,000 times. Cophenetic correlations and Baker’s gamma coefficients between dendrograms were computed using the cor_cophenetic and cor_bakers_gamma functions of dendextend, respectively, with significance again evaluated via permutation tests (1,000 shuffles).

#### Sensitivity analysis for strain-level baseline effects

To assess the potential impact of strain-level baseline growth differences on interstrain correlations, we performed a sensitivity analysis using strain-baseline-normalized growth-rate profiles. Briefly, growth rates were z-scored within each carbon source across strains, and each strain’s mean z-score was then subtracted; interstrain correlations were recomputed from these normalized profiles and the resulting dendrogram was compared with the original (**Figure S6**).

#### Sensitivity analysis for zero-growth entries

To assess the potential impact of zero-growth entries on interstrain correlations, we first quantified the frequency of strain–substrate pairs involving at least one zero-growth condition within each cluster (**Figure S7A**). We then repeated the correlation analysis and hierarchical clustering after excluding all zero-growth samples (**Figure S7B–D**). As a more stringent approach, strains exhibiting zero growth on any of the retained carbon sources were additionally excluded, reducing the dataset from 70 to 53 strains and from 1,732 to 1,325 samples; hierarchical clustering was repeated on this filtered dataset (**Figure S7E–G**). Dendrogram topology was compared between the original and filtered datasets using entanglement scores and cophenetic correlations with permutation tests (1,000 shuffles), as described above.

#### Principal component analysis (PCA)

PCA was conducted using the 22 carbon sources shared between the ECOR and MA datasets. Growth-rate profiles of the MA lineages were normalized and subjected to PCA using the prcomp function in R. ECOR growth-rate profiles were normalized in the same manner and projected onto the MA-derived PC space by multiplying them with the MA eigenvectors. The resulting PC scores for each ECOR strain were used to calculate cumulative explained variance.

To assess whether observed variance structure differed from that expected under random conditions, we created 1,000 randomly label-shuffled datasets derived from the ECOR matrix (70 × 22) using the sample function in R. Each dataset was projected onto the MA-derived PC space to compute the distribution of cumulative explained variance, enabling estimation of the mean and 95% confidence intervals for the null expectation.

