## Supplemental Figures for "Low-Dimensional Organization of Metabolic Phenotypes Underlies Recurrent Evolutionary Diversification"

### Supplementary information for
### Low-Dimensional Organization of Metabolic Phenotypes Underlies Recurrent Evolutionary Diversification

Keita Saito[1,2], Saburo Tsuru[3,*], and Chikara Furusawa[2,3,4,*]

[1]Graduate School of Science and Engineering, Chiba University, 1-33 Yayoi-cho, Inage-ku, Chiba-city, Chiba, 263-8522, Japan
[2]Center for Biosystems Dynamics Research (BDR), RIKEN, 6-7-1 Minatojima-minamimachi, Chuo-ku, Kobe, 650-0047, Japan
[3]Universal Biology Institute, Graduate School of Science, The University of Tokyo, 7-3-1 Hongo, Bunkyo, Tokyo 113-0033, Japan
[4]Department of Physics, Graduate School of Science, The University of Tokyo, 7-3-1 Hongo, Bunkyo, Tokyo 113-0033, Japan

*Corresponding Authors
Saburo Tsuru

Chikara Furusawa


Supplementary Figures

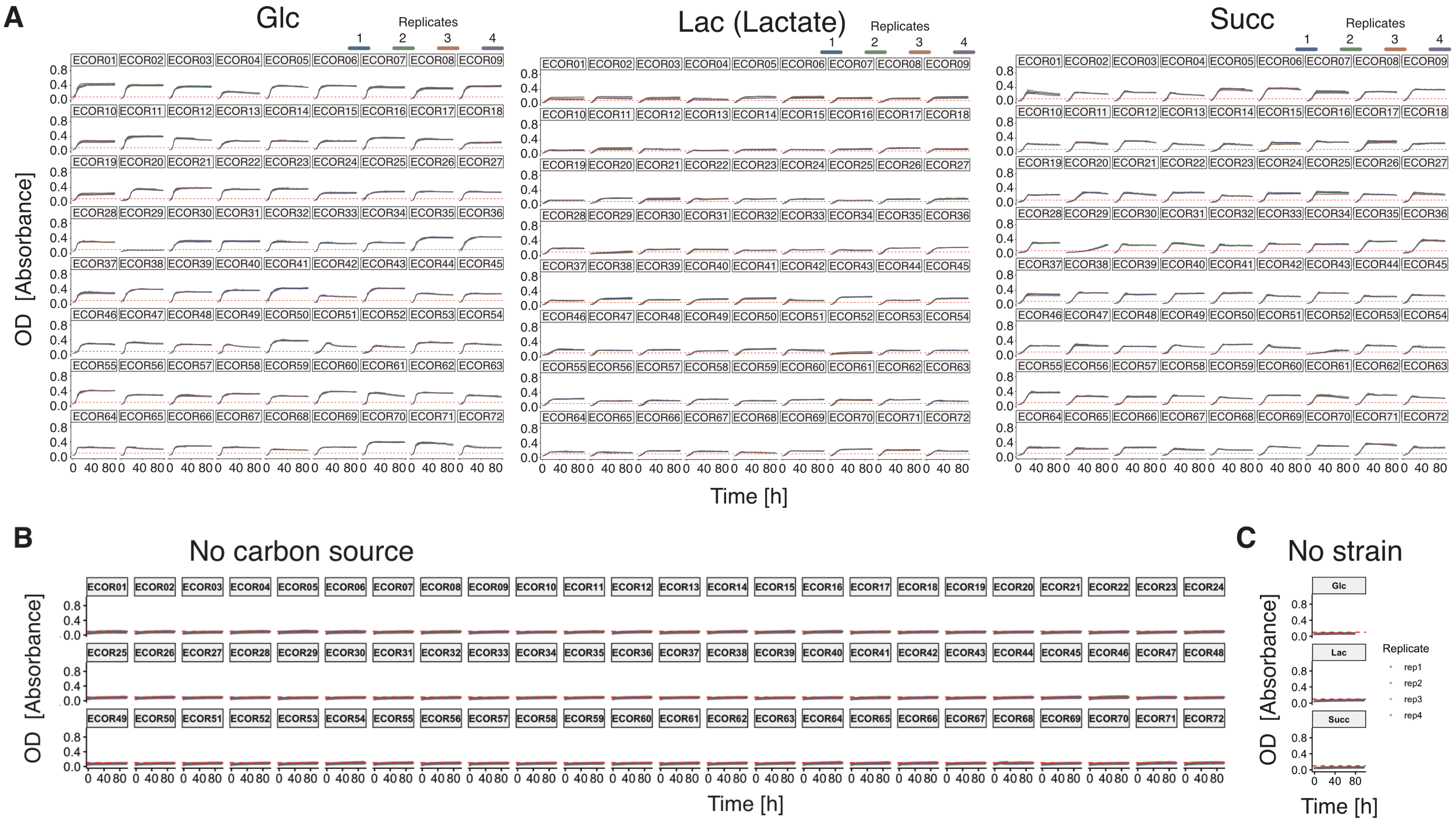


**Figure S1 Representative growth curves.** (**A**) Growth curves for 72 ECOR strains are shown for representative carbon sources from conventional metabolic categories: glucose (Glc) for upper glycolysis, lactate (Lac) for lower glycolysis, and succinate (Succ) for the TCA cycle. Gray points indicate experimental data from four replicates, while colored lines represent fits to a logistic model. (**B**) Temporal changes in OD of cultures without carbon sources. (**C**) Temporal changes in OD of sterile cultures. Horizontal dashed lines represent the threshold (OD of 0.1) used for growth categorization.

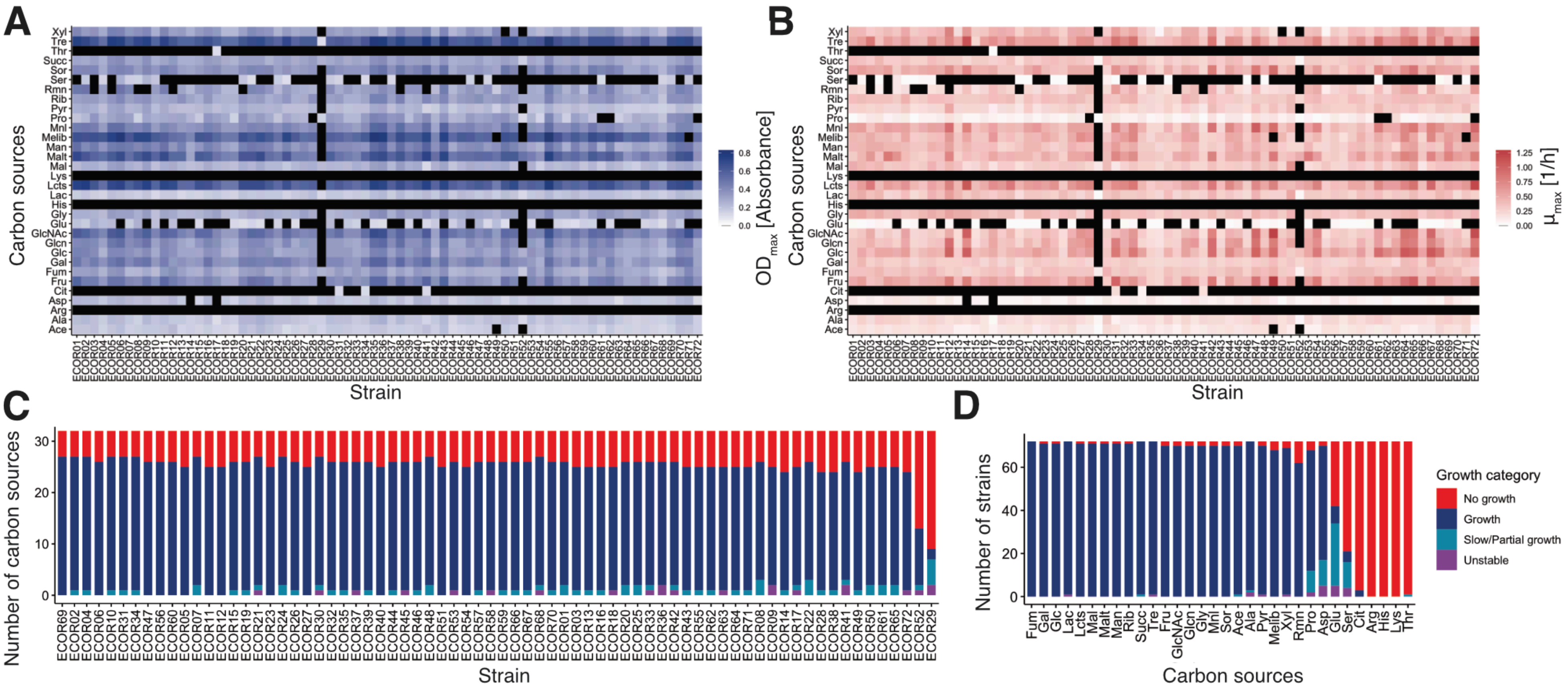


**Figure S2 Growth characteristics of ECOR strains.** (**A**) and (**B**) Heat maps of the mean $OD_{max}$ (**A**) and $\mu_{max}$ (**B**) across 32 carbon sources are shown. (**C**) and (**D**) Stacked bar charts representing the distribution of growth categories across strains (**C**) and carbon sources (**D**). In both panels, the x-axis is sorted in descending order (left to right) based on the frequency of the "Growth" category. Lac and Lcts represent lactate and lactose, respectively.

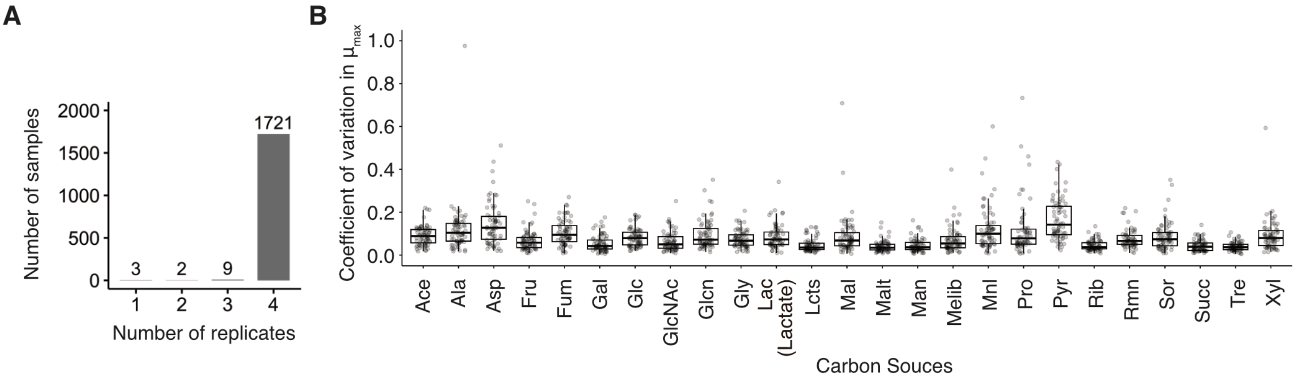


**Figure S3 Number of replicates and replicate-based coefficient of variation (CV) of growth rates across carbon sources in ECOR.** (**A**) Number of replicates used to calculate the mean, variance, and coefficient of variation (CV) of the maximum growth rate ($\mu_{max}$) for each strain–carbon source combination. (**B**) Distribution of the coefficient of variation (CV) of growth rates across replicates for each carbon source. Individual data points (jitters) represent individual strains. Boxplot elements represent the median (horizontal line), first and third quartiles (q1 and q3), and whiskers extending to the most extreme data points within 1.5×(q3 − q1). Lac and Lcts represent lactate and lactose, respectively.

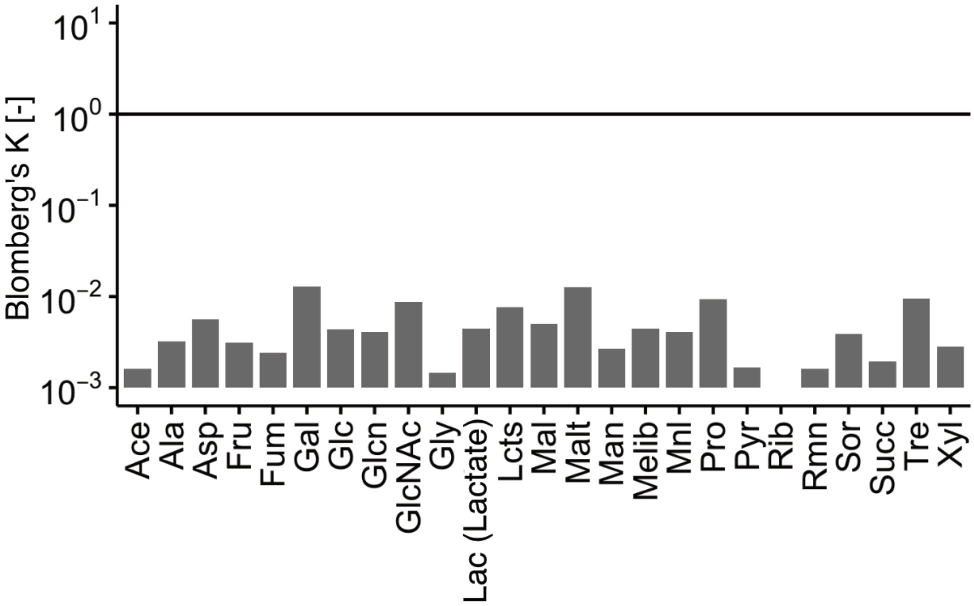


**Figure S4 Phylogenetic signal in growth rates of ECOR strains.** The horizontal line represents a Blomberg's *K* value of 1, which indicates that the observed trait distribution perfectly matches the expectation under a Brownian motion model of evolution. Observed *K* values were substantially lower than 1 across all conditions, indicating a very weak phylogenetic signal. Furthermore, BH-adjusted P-values exceeded 0.05 for all conditions, confirming the lack of significant phylogenetic signal. Lac and Lcts represent lactate and lactose, respectively.

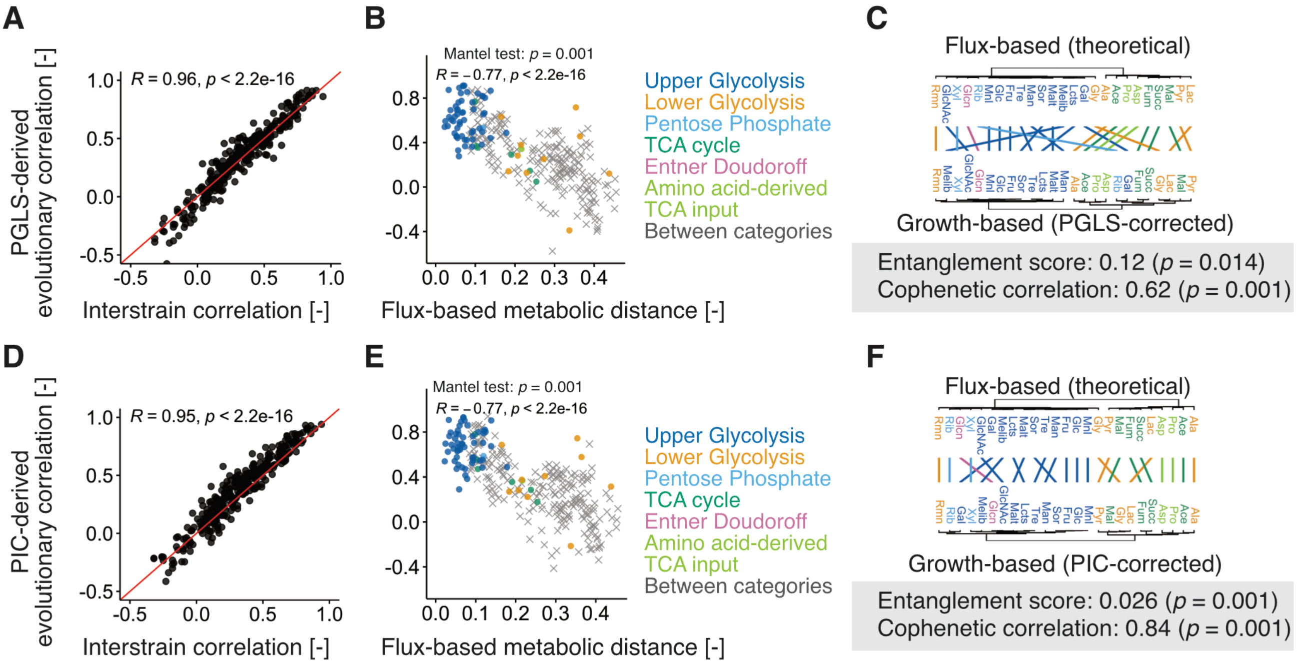


**Figure S5 Comparison between uncorrected interstrain correlations and phylogeny-corrected evolutionary correlations.** Panels (**A**)–(**C**) and (**D**)–(**F**) show results based on Phylogenetic Generalized Least Squares (PGLS) and Phylogenetically Independent Contrasts (PIC), respectively. (**A**) and (**D**) Relationship between uncorrected interstrain correlations and PGLS-derived (**A**) or PIC-derived (**D**) evolutionary correlations. The solid line represents y = x. (**B**) and (**E**) Relationship between flux-based metabolic distance and PGLS-derived (**B**) or PIC-derived evolutionary correlations. Spearman's R and associated two-sided P-values, as well as the P-values from Mantel tests with 999 permutations, are indicated. Symbols denote within- (circles) and between-category (crosses) pairs. (**C**) and (**F**) Tanglegrams comparing the flux-based dendrogram (top) and the growth-based dendrograms derived from PGLS-derived (**C**) or PIC-derived (**F**) correlations (bottom). All dendrograms were constructed by hierarchical clustering (Ward.D2). The entanglement score and cophenetic correlation are indicated. Lac and Lcts represent lactate and lactose, respectively.

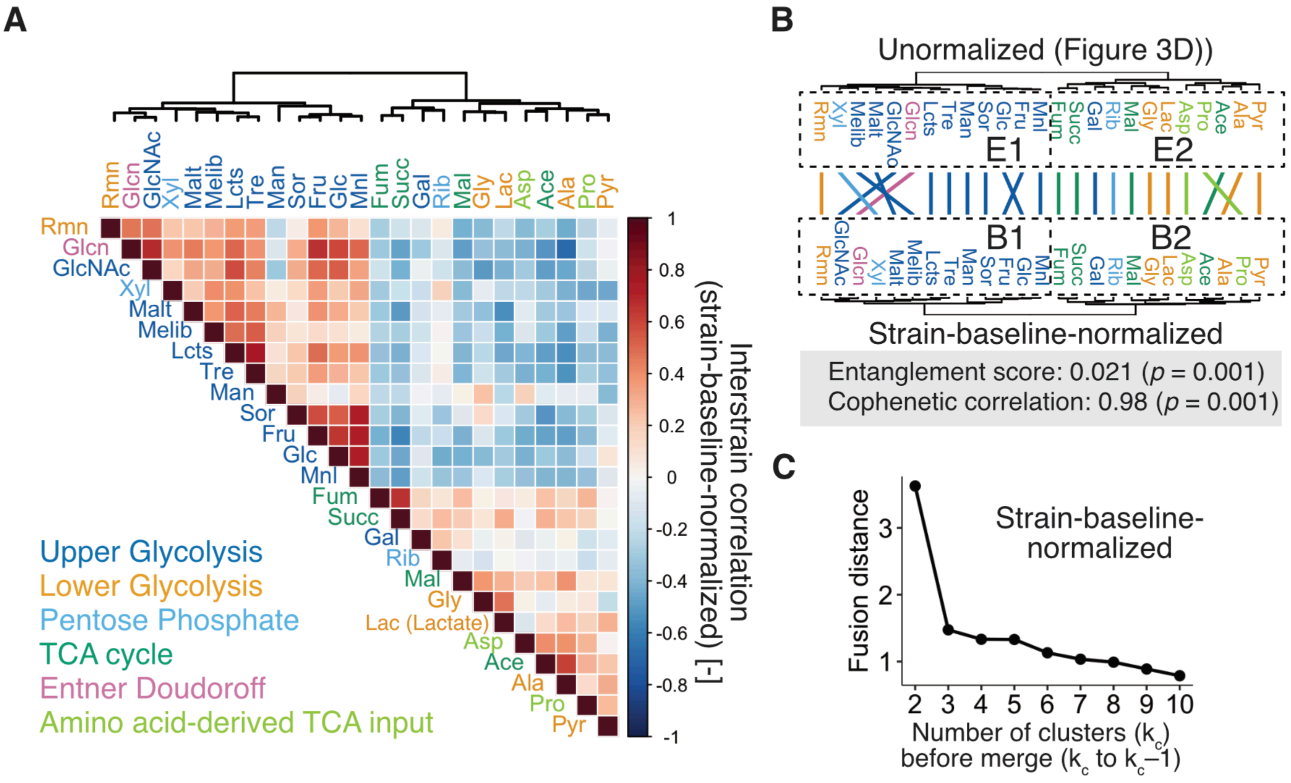


**Figure S6 Interstrain correlations using strain-baseline-normalized growth-rate profiles in ECOR.** (**A**) Heatmap of interstrain correlations among all carbon-source pairs calculated from strain-baseline-normalized growth rates (after subtracting each strain's mean growth rate across all carbon sources). Lac and Lcts represent lactate and lactose, respectively. (**B**) Tanglegram comparing the original unnormalized dendrogram (top, **Figure 3D**) and the dendrogram derived from the baseline-normalized data (bottom). Both dendrograms were constructed by hierarchical clustering (Ward.D2). The entanglement score and cophenetic correlation are indicated. Dashed boxes represent clusters based on hierarchical clustering (clusters B1 and B2). (**C**) Scree plot of the dendrogram's fusion distances across different numbers of clusters ($k_c$ prior to merging to $k_c - 1$).

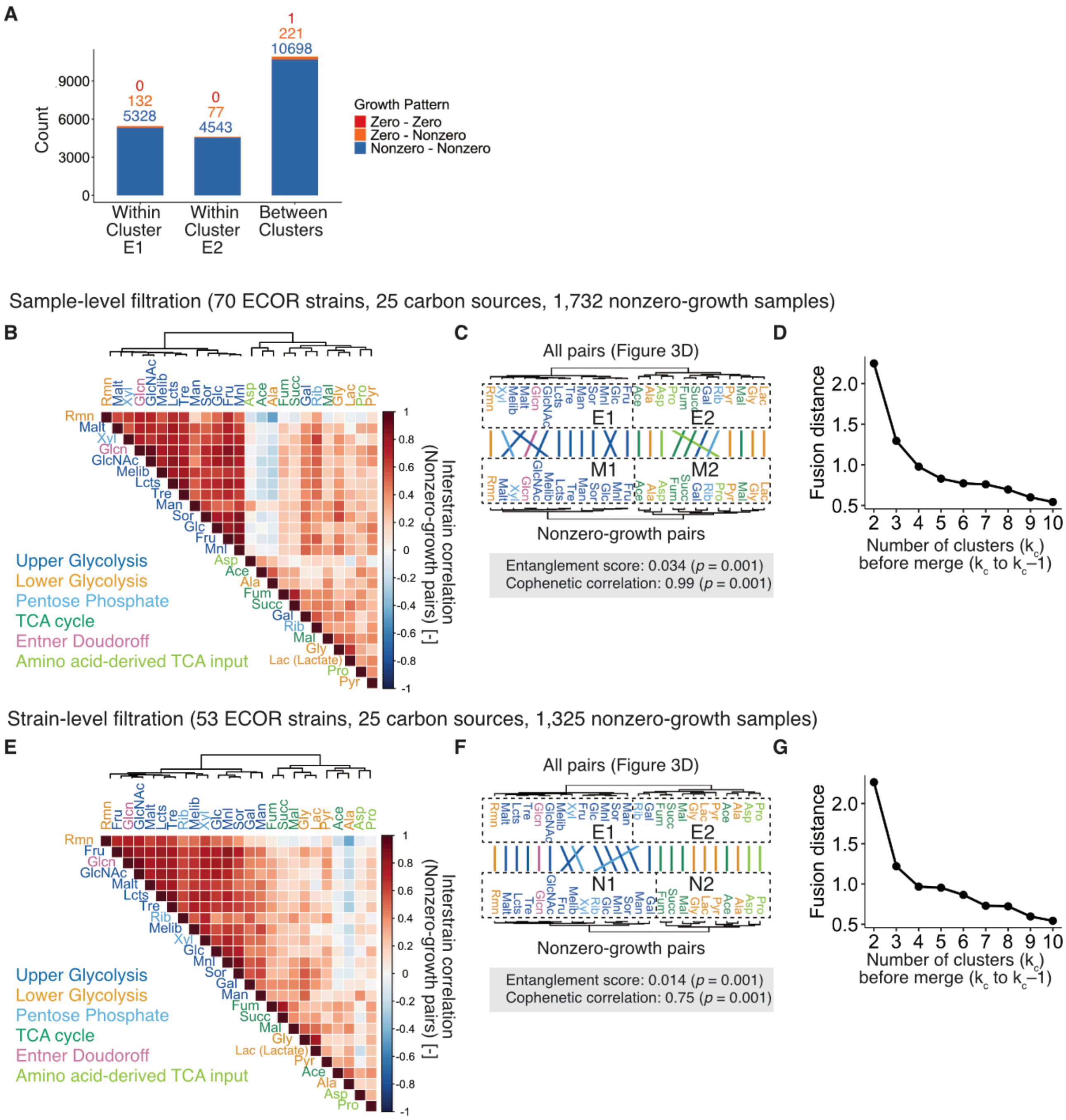


**Figure S7 Evaluation of interstrain correlations excluding zero-growth data in ECOR.** (**A**) Distribution of growth patterns (Zero–Zero, Zero–Nonzero, and Nonzero–Nonzero) across strain-carbon pairs within and between clusters, demonstrating that zero-growth conditions are extremely rare. (**B**)–(**D**) Analysis of sample-level filtration. Eighteen zero-growth samples were removed, yielding 1,732 nonzero-growth samples with 70 ECOR strains and 25 carbon sources. (**E**)–(**G**) Analysis of strain-level filtration. Strains containing zero-growth samples were removed, yielding 1,325 nonzero-growth samples with 53 ECOR strains and 25 carbon sources. (**B**) and (**E**) Heatmap of interstrain correlations among all carbon-source pairs calculated exclusively from nonzero-growth data (**B** for sample-level and **E** for strain-level filtration). (**C**) and (**F**) Tanglegrams comparing the original dendrograms (top; **Figure 3D**) and the filtered dendrogram derived from nonzero-growth pairs (bottom; **C** for sample-level and **F** for strain-level filtration). All dendrograms were constructed by hierarchical clustering (Ward.D2). The entanglement scores and cophenetic correlations are indicated. Dashed boxes represent clusters based on hierarchical clustering (clusters M1 and M2 in **C**; clusters N1 and N2 in **F**). (**D**) and (**G**) Scree plots of the dendrogram's fusion distances across different numbers of clusters ($k_c$ prior to merging to $k_c - 1$; (**D**) for sample-level and (**G**) for strain-level filtration). Lac and Lcts represent lactate and lactose, respectively.

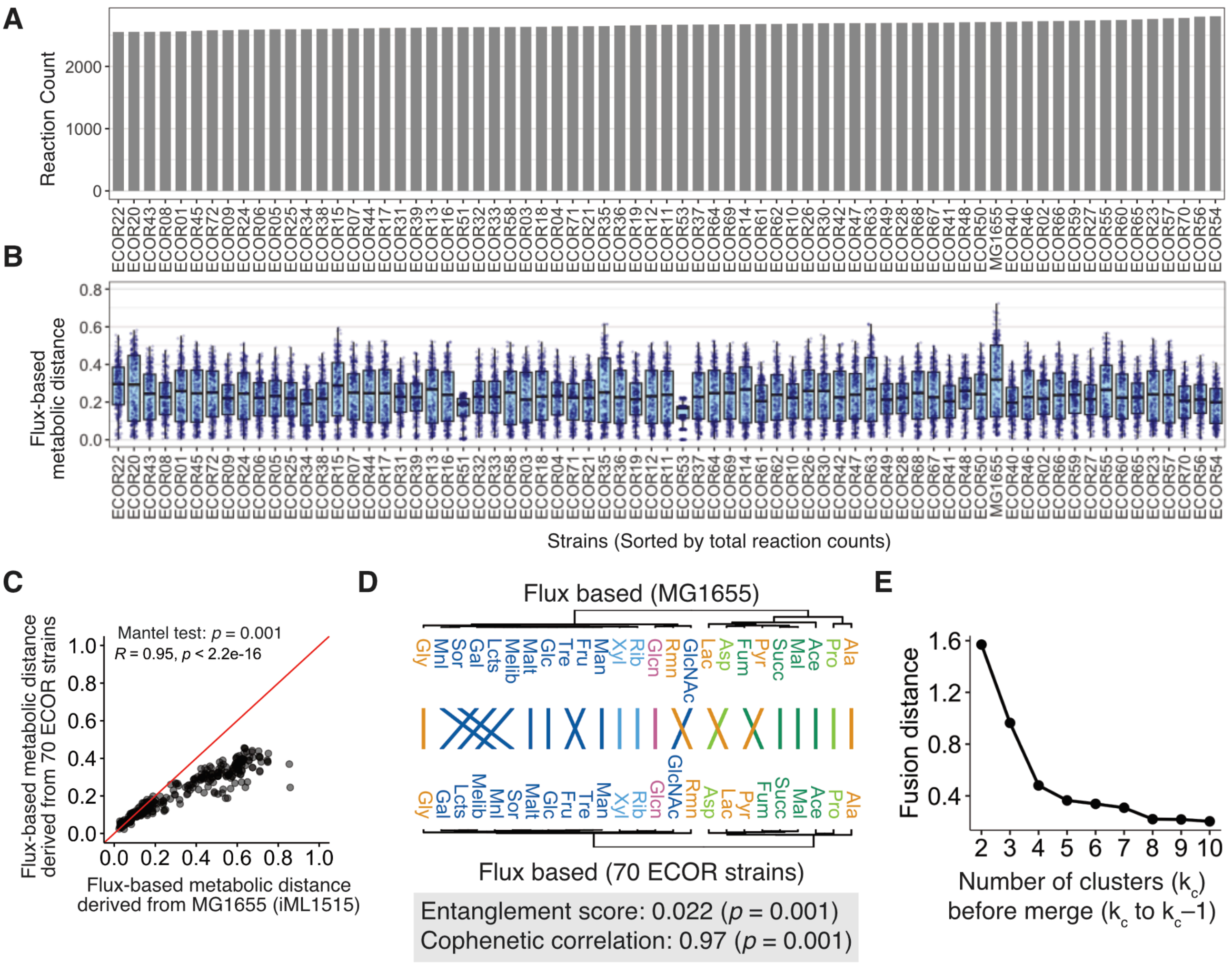


**Figure S8 Flux-based metabolic distances across genome-scale metabolic models.** (**A**) Total reaction counts for each of the 70 ECOR models and iML1515 (MG1655), sorted in ascending order (from left to right). (**B**) Distribution of flux-based metabolic distances across models. Individual data points (jitters) represent different carbon source pairs. Boxplot conventions are identical to those in **Figure S3B**. (**C**) Comparison of flux-based metabolic distances derived from the 70 ECOR strains (mean) with those derived from the reference model (iML1515 for K-12 MG1655). Each point represents a different pair of carbon sources. The solid line represents y = x. (**D**) Tanglegram comparing the flux-based dendrogram derived from the iML1515 model (top) and the dendrogram derived from the 70 ECOR strains (bottom). Both dendrograms were constructed by hierarchical clustering (Ward.D2). The entanglement score and cophenetic correlation are indicated. (**E**) Scree plot of the dendrogram's fusion distances across different numbers of clusters ($k_c$ prior to merging to $k_c - 1$) for the iML1515 model. Lac and Lcts represent lactate and lactose, respectively.

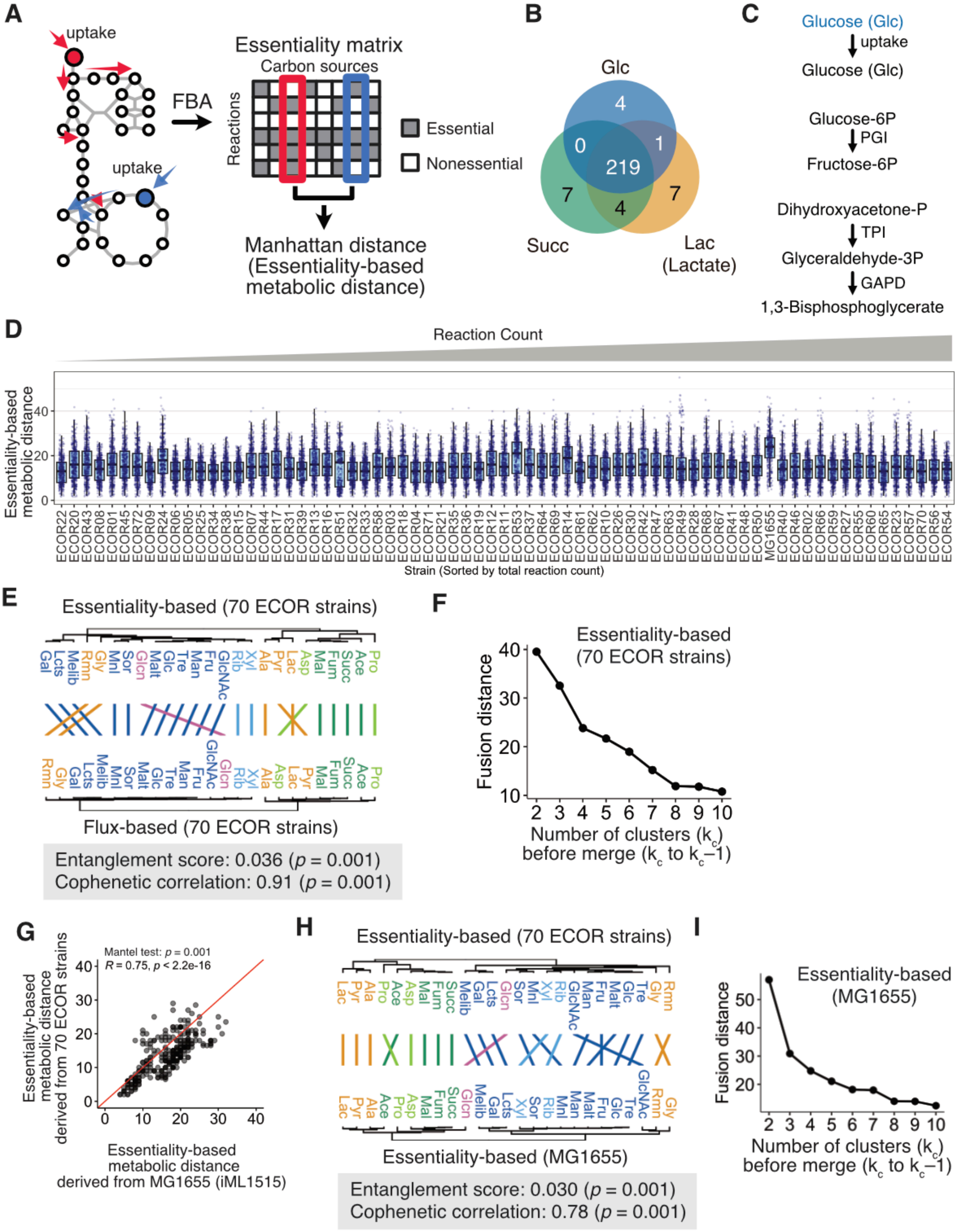


**Figure S9 Essentiality-based metabolic distances across genome-scale metabolic models.** (A) Schematic representation of essentiality-based metabolic distance calculation. FBA with single-reaction knockouts using genome-scale metabolic models identified essential reactions for optimal biomass production. A reaction was classified as essential if its deletion decreased optimal biomass flux by more than 0.1%. The Manhattan distance between essential-reaction vectors was used as the essentiality-based metabolic distance. (**B**) Representative Venn diagram showing the overlap of essential reactions among three carbon sources in ECOR 11. (**C**) The four essential reactions unique to glucose, as identified in panel (**B**). (**D**) Distribution of essentiality-based metabolic distances across models. Individual data points (jitters) represent different carbon source pairs. Boxplot conventions are identical to those in **Figure S3B**. Models are sorted by total reaction counts in ascending order (from left to right). (**E**) Tanglegram comparing the essentiality-based (top) and flux-based (bottom) dendrograms. Both dendrograms were constructed from the 70 ECOR strains. (**F**) Scree plot of the dendrogram's fusion distances across different numbers of clusters ($k_c$ prior to merging to $k_c - 1$) for the 70 ECOR strains. (**G**) Comparison of essentiality-based metabolic distances derived from the 70 ECOR strains with those derived from the reference model (iML1515 for K-12 MG1655). Each point represents a different pair of carbon sources. The solid line represents $y = x$. (**H**) Tanglegram comparing the essentiality-based dendrogram derived from the 70 ECOR models (top) and the dendrogram derived from the iML1515 model (bottom). In panels (**E**) and (**H**), all dendrograms were constructed by hierarchical clustering (Ward.D2). The entanglement scores and cophenetic correlations are indicated. (**I**) Scree plot of the dendrogram's fusion distances across different numbers of clusters ($k_c$ prior to merging to $k_c - 1$) for the iML1515 model (MG1655). Lac and Lcts represent lactate and lactose, respectively.

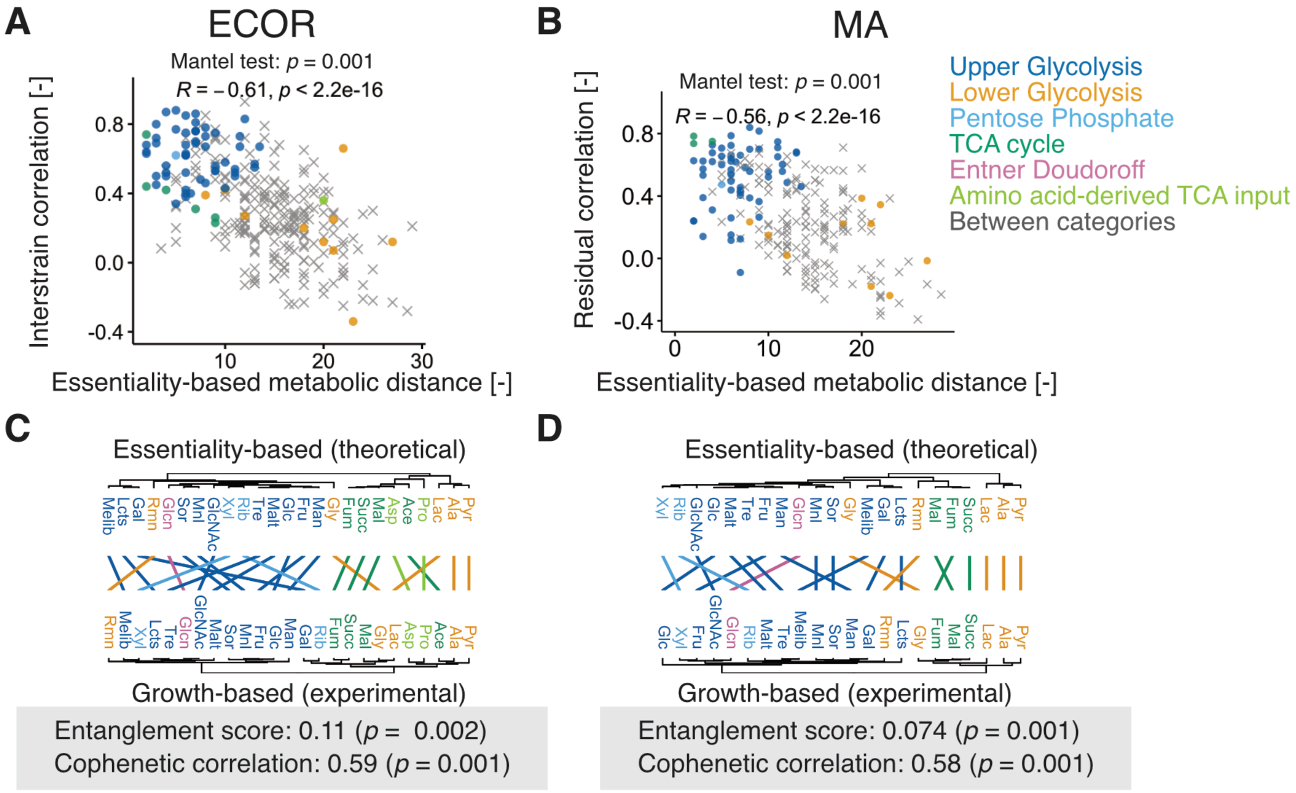


**Figure S10 Correlation analysis using essentiality-based metabolic distance.** (**A**) and (**B**) Relationship between essentiality-based metabolic distance and interstrain (**A**) or residual (**B**) correlation. Spearman's R and associated two-sided P-values, as well as the P-values from Mantel tests with 999 permutations, are indicated. Symbols denote within- (circles) and between-category (crosses) pairs. (**C**) and (**D**) Tanglegrams comparing the essentiality-based dendrogram (top) and the dendrogram derived from interstrain correlation in ECOR (**C**, bottom) or residual correlation in MA (**D**, bottom). Both dendrograms were constructed by hierarchical clustering (Ward.D2). The entanglement scores and cophenetic correlations are indicated. Lac and Lcts represent lactate and lactose, respectively.

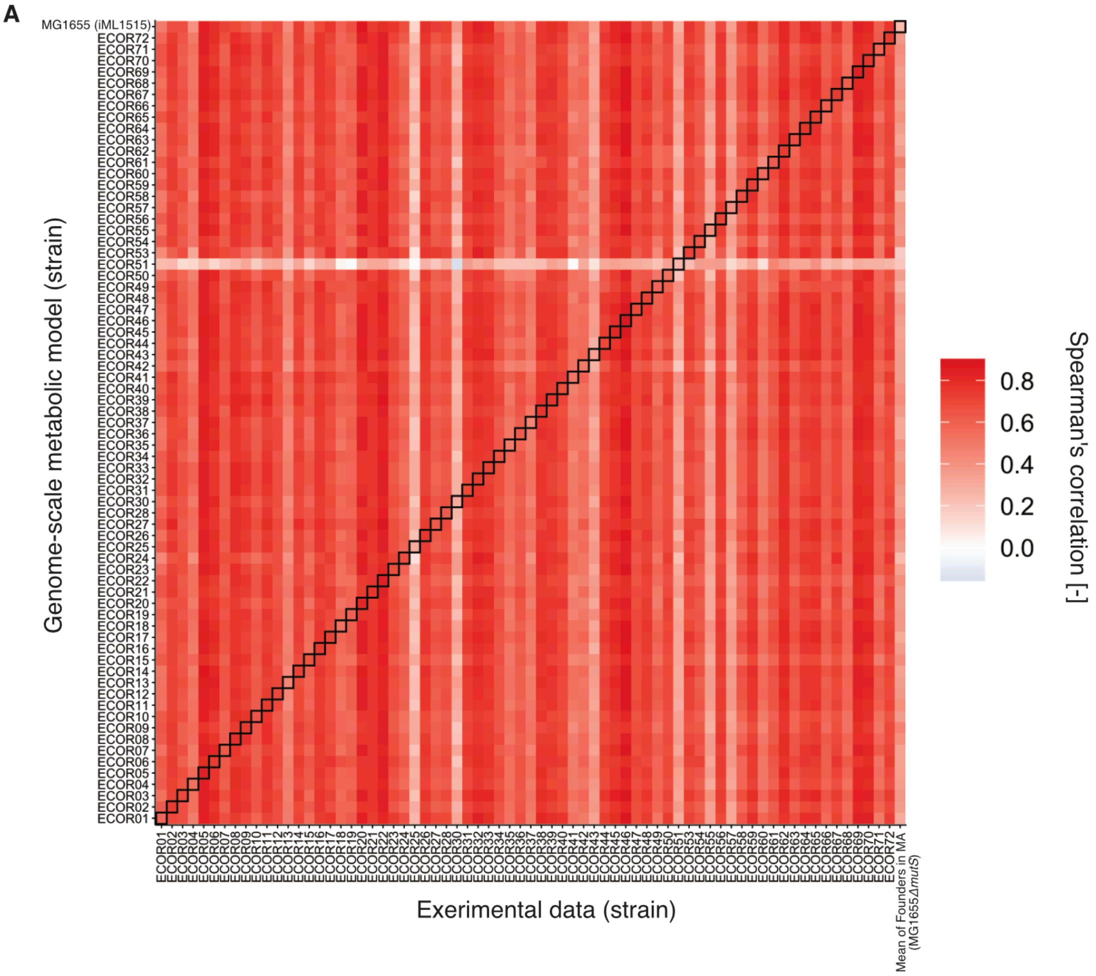


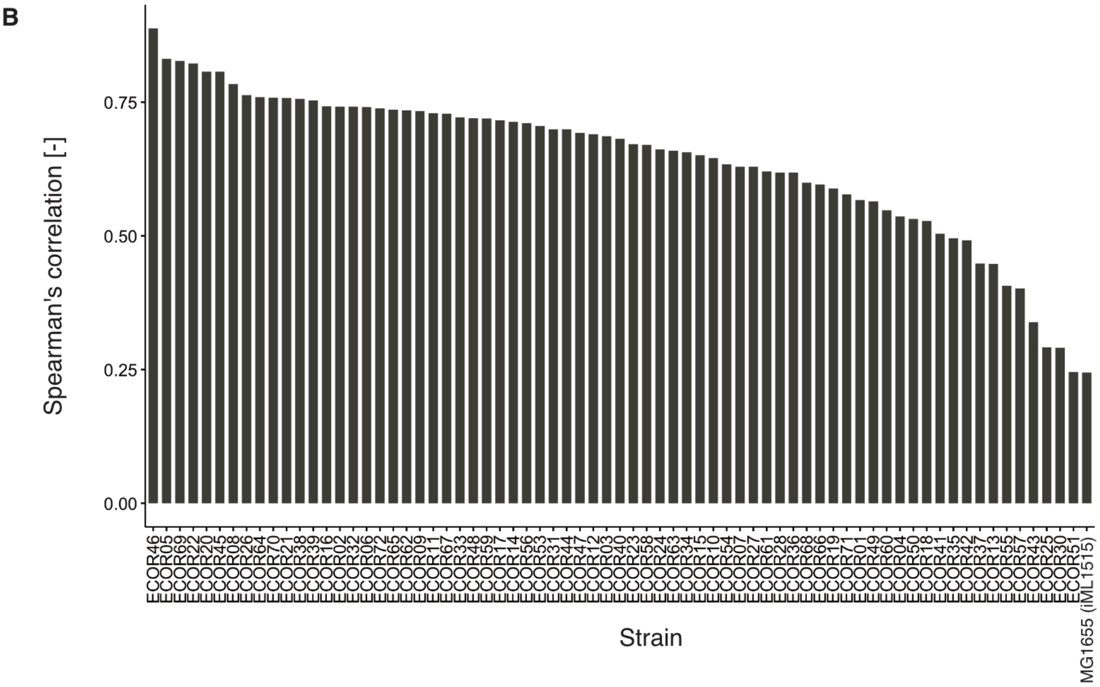


**Figure S11 Cross-strain FBA prediction performance.** (**A**) Heatmap displaying Spearman's rank correlation coefficients between experimentally measured growth rates ($\mu_{max}$) across carbon sources and Flux Balance Analysis (FBA) predictions (biomass production rate) for all pairwise combinations of FBA models and strain experimental data. Columns represent experimental data for each strain, while rows correspond to genome-scale metabolic models. Cells outlined with black borders highlight matching model–strain pairs (self-prediction, where the model strain equals the experimental strain). The mean growth rate of all three founders in the MA experiment was used as a reference growth rate for MG1655 $\Delta$*mutS*, which is assumed to be the strain in this study closest to MG1655, modeled using iML1515. (**B**) Spearman's correlation coefficients for self-prediction across strains, corresponding directly to the black-outlined cells in panel (**A**). Each bar represents the predictive accuracy of a strain-specific FBA model when evaluated against its own experimental growth data across carbon sources.

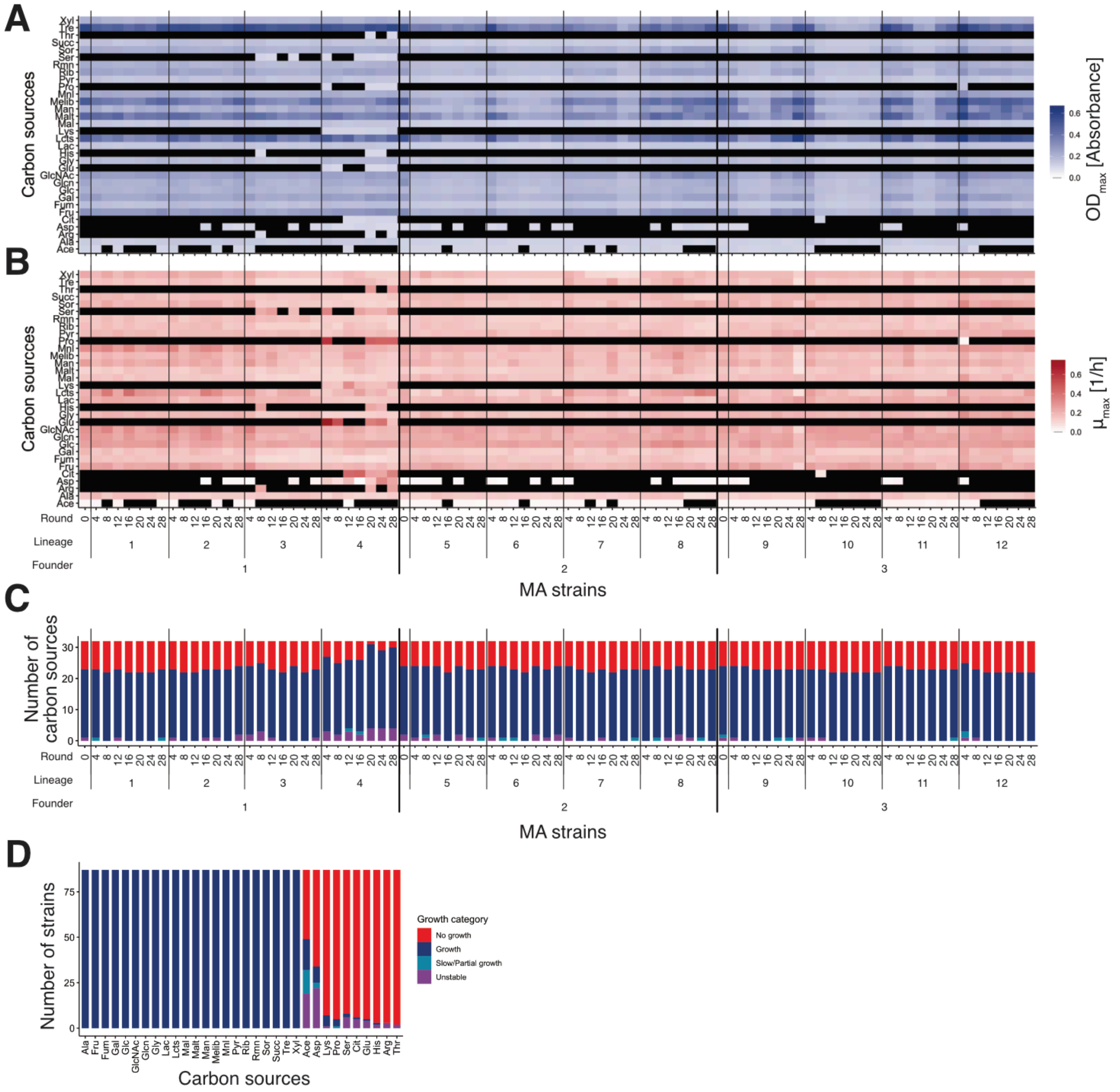


**Figure S12 Growth characteristics of MA lineages.** (**A**) and (**B**) Heat maps showing the mean $OD_{max}$ (**A**) and $\mu_{max}$ (**B**) across carbon sources. (**C**) and (**D**) Stacked bar charts representing the distribution of growth categories across lineages (**C**) and carbon sources (**D**). In both panels, the x-axis is sorted in descending order (left to right) based on the frequency of the "Growth" category. Lac and Lcts represent lactate and lactose, respectively.

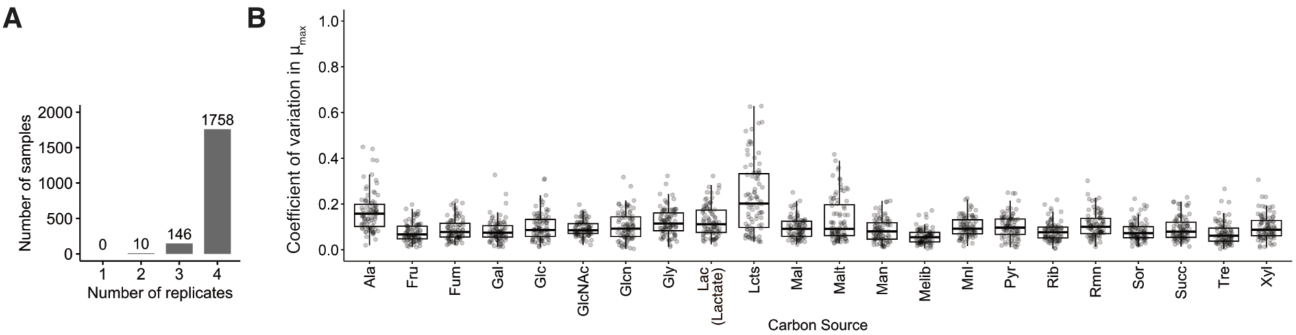


**Figure S13 Number of replicates and replicate-based CV of growth rates across carbon sources in MA.** (**A**) Number of replicates used to calculate the mean, variance, and CV of the maximum growth rate ($\mu_{max}$) for each sample–carbon source combination. (**B**) Distribution of the CV of growth rates across replicates for each carbon source. Boxplot conventions are identical to those in **Figure S3B**. Lac and Lcts represent lactate and lactose, respectively.

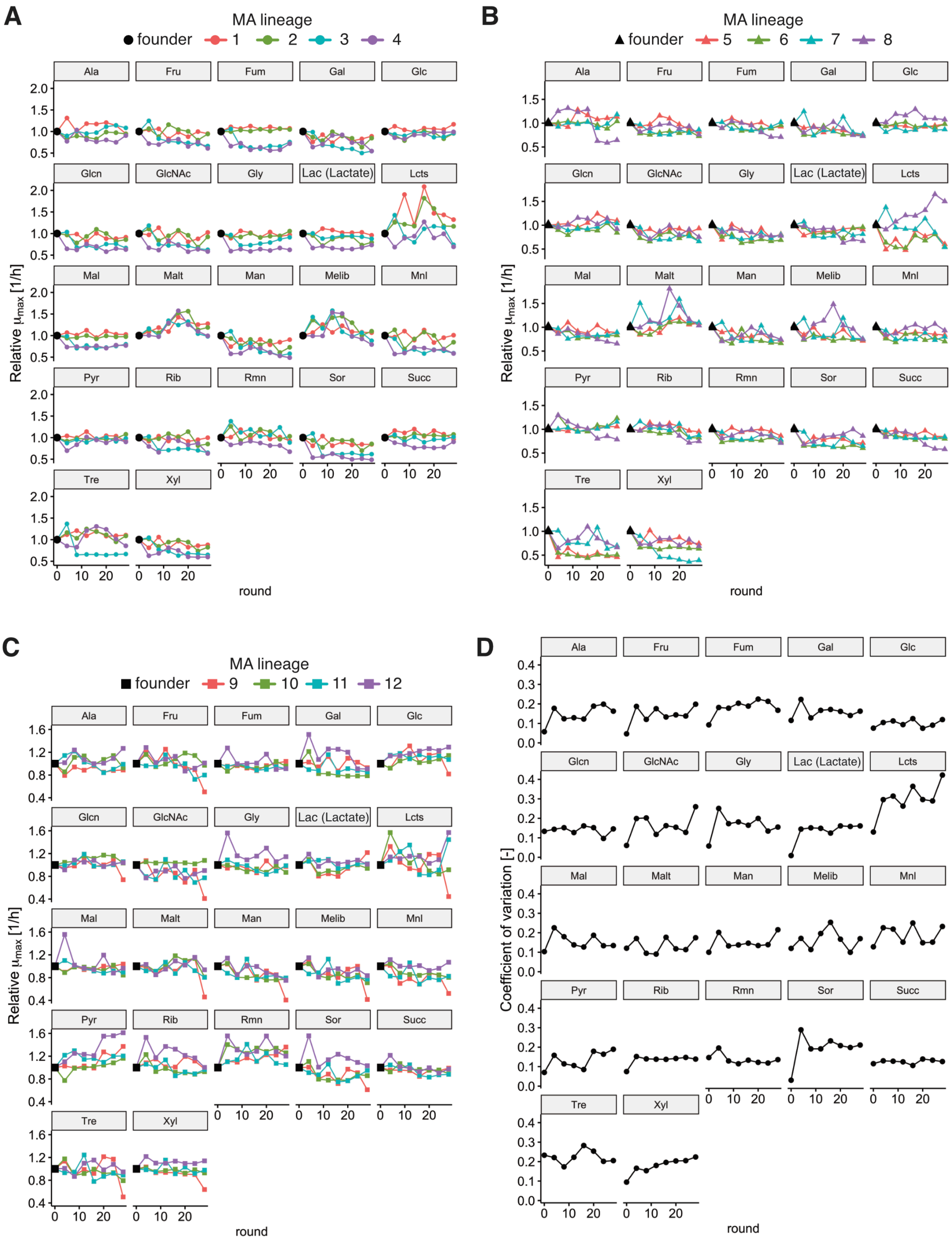


**Figure S14 Dynamics of growth rates during MA experiments.** (**A**)–(**C**) Changes in relative growth rates across successive MA rounds for lineages derived from founder 1 (**A**), founder 2 (**B**), and founder 3 (**C**). The relative growth rates were defined as growth rates normalized by dividing by the corresponding founder values. This normalization was used only for visualization and was not used in the LMM analysis. (**D**) The coefficient of variation (CV; defined as the standard deviation divided by the mean) calculated from raw growth rates across lineages over the course of MA rounds. Lac and Lcts represent lactate and lactose, respectively.
